# Central Role for Glycolysis and Fatty Acids in LH-responsive Progesterone Synthesis

**DOI:** 10.1101/2024.02.14.580329

**Authors:** Emilia Przygrodzka, Fatema Binderwala, Robert Powers, Renee M. McFee, Andrea S. Cupp, Jennifer R. Wood, John S. Davis

## Abstract

Progesterone production by the corpus luteum is fundamental for establishing and maintaining pregnancy. The pituitary gonadotropin luteinizing hormone (LH) is recognized as the primary stimulus for luteal formation and progesterone synthesis, regardless of species. Previous studies demonstrated an elevation in abundance of genes related to glucose and lipid metabolism during the follicular to luteal transition. However, the metabolic phenotype of these highly steroidogenic cells has not been studied. Herein, we determined acute metabolic changes induced by LH in primary luteal cells and defined pathways required for progesterone synthesis. Untargeted metabolomics analysis revealed that LH induces rapid changes in vital metabolic pathways, including glycolysis, tricarboxylic acid (TCA) cycle, pentose phosphate pathway, *de novo* lipogenesis, and hydrolysis of phospholipids. LH stimulated glucose uptake, enhanced glycolysis, and flux of [U-^13^C_6_]-labeled glucose-derived carbons into metabolic branches associated with adenosine 5’-triphosphate (ATP) and NADH/NADPH production, synthesis of nucleotides, proteins, and lipids, glycosylation of proteins or lipids, and redox homeostasis. Selective use of small molecule inhibitors targeting the most significantly changed pathways, such as glycolysis, TCA cycle, and lipogenesis, uncovered cellular metabolic routes required for LH-stimulated steroidogenesis. Furthermore, LH via the protein kinase A (PKA) pathway triggered *post-*translational modification of acetyl-CoA carboxylase alpha (ACACA) and ATP citrate lyase (ACLY), enzymes involved in *de novo* synthesis of fatty acids. Inhibition of ACLY and fatty acid transport into mitochondria reduced LH-stimulated ATP, cAMP production, PKA activation, and progesterone synthesis. Taken together, these findings reveal novel hormone-sensitive metabolic pathways essential for maintaining LHCGR/PKA signaling and steroidogenesis in ovarian luteal cells.

**Significance:** The establishment and maintenance of pregnancy require a well-developed corpus luteum, an endocrine gland within the ovary that produces progesterone. Although there is increased awareness of intracellular signaling events initiating the massive production of progesterone during the reproductive cycle and pregnancy, there are critical gaps in our knowledge of the metabolic and lipidomic pathways required for initiating and maintaining luteal progesterone synthesis. Here, we describe rapid, hormonally triggered metabolic pathways, and define metabolic targets crucial for progesterone synthesis by ovarian steroidogenic cells. Understanding hormonal control of metabolic pathways may help elucidate approaches for improving ovarian function and successful reproduction or identifying metabolic targets for developing nonhormonal contraceptives.

## 1. Introduction

The mammalian ovary has two principal functions: the production and release of mature ova capable of fertilization, and the synthesis and secretion of steroids, which are critical to ensure ovulation and pregnancy, but additionally to impact secondary sex characteristics, as well as bone, brain, immune, and cardiovascular health and metabolism (1–3). Within the ovary, ovarian follicles consisting of theca and granulosa cells provide a developmental niche allowing appropriate maturation of ova (4, 5). Mature ovarian follicles rupture during ovulation releasing ova into the fallopian tube, while the remnant of the follicle transforms into the corpus luteum, a highly vascularized, progesterone secreting powerhouse (3, 6). Progesterone, a steroid hormone produced by this transient endocrine gland is essential for ovulation (7), transport of ova through the fallopian tube (8), embryonic development, establishment and maintenance of the uterine environment to initiate and sustain pregnancy as well as development of mothering behavior (9, 10). Recently, it was found that presence of the corpus luteum reduces the risk of preeclampsia, the most common pregnancy complication in IVF patients (11). If fertilization does not occur, the corpus luteum regresses and the reduction in progesterone allows for the progression of a new menstrual or estrous cycle (3, 12). Inadequate progesterone production at critical periods can lead to premature pregnancy loss in women and domestic animals (13). Given the importance of progesterone to successful reproduction, and increasing infertility rates, studies to understand the hormonal and cellular control of progesterone offer insight into fertility regulation.

Among various luteotrophic factors, luteinizing hormone (LH), a gonadotropin released by the pituitary gland, plays a crucial role in the formation, development, and maintenance of the corpus luteum in women and mammalian animals (3, 6). LH binds to its cognate receptor (LHCGR) present on the surface of luteal cells and rapidly elevates the production of cyclic adenosine monophosphate (cAMP), which consequently activates protein kinase A (PKA) (14–16). At the time of ovulation, this cascade of events recruits additional signaling events that in combination alter the phosphorylation of transcription factors enhancing the expression of genes associated with angiogenesis, ovulation, cellular differentiation, steroidogenesis, and cell survival (17–20). These events are also responsible for the post-transcriptional regulation of proteins involved in the maintenance of luteal function like enhanced lipolysis required to sustain cholesterol supply for progesterone synthesis or changes in mitochondrial dynamics (21–23). Considerable effort has been placed in the past five decades to understand how gonadotropins control ovarian gene transcription and post-translational modifications of various signaling proteins that positively or negatively impact transcription (21, 22, 24, 25). Less is known regarding the metabolic events driven by LH that contribute to progesterone secretion by the corpus luteum.

Our current understanding of the metabolic phenotype of luteal cells is limited and based largely on the luteal transcriptome and proteome (18, 26). Recently, we identified changes in the expression of genes encoding proteins associated with the metabolism of glucose and lipids in luteal cells in comparison to ovarian follicle cells (18, 27). However, significant gaps in our knowledge exist concerning the metabolic pathways governing the trophic actions of LH on steroidogenesis and maintenance of luteal function. Metabolic profiling of steroidogenic luteal cells can provide insight into the cellular metabolic pathways that support the pronounced increase in steroidogenesis observed following ovulation. Herein, using two unbiased metabolic approaches, i.e., mass spectrometry and nuclear magnetic resonance (NMR), we characterized the temporal metabolic changes in highly LH-responsive steroidogenic luteal cells. Further analysis determined the effects of LH on mitochondrial respiration, glucose uptake and glycolysis. We also evaluated the significance of specific LH-mediated metabolic pathways controlling steroidogenesis by using selective small molecule inhibitors. We identified acetyl-CoA carboxylase alpha (ACACA) and ATP citrate lyase (ACLY), two enzymes involved in *de novo* lipogenesis, as targets for LH-stimulated PKA signaling, and disruption of ACLY phosphorylation or enzyme activity as well as transport of fatty acids to mitochondria acutely blocked LHCGR/PKA signaling and LH-stimulated progesterone synthesis. Notably, this study was conducted with LH-responsive primary steroidogenic small luteal cells from the bovine corpus luteum, which is powerful model to study human ovarian physiology (28). Like women, cows are mono-ovulatory with multiple waves of follicular development, and the length of the luteal phase and pregnancy are similar (28). Moreover, the signaling pathways triggered by LHCGR are evolutionary conserved (3) and LH is necessary for the formation and maintenance of the corpus luteum, and progesterone is required to establish and maintain pregnancy in both species (28).

## 2. Results

### LH induces acute metabolic alterations in steroidogenic luteal cells

This study was designed to determine the rapid, non-genomic action of LH on acute metabolic changes in steroidogenic bovine small luteal cells. To accomplish this, an untargeted approach was used to measure metabolites in the whole cell extracts and conditioned medium of luteal cells after incubation with LH (10 ng/ml) for 10, 30, 60, and 240 min (Figure 1A-F; Supplementary Table 1). Because luteal cells continuously produce progesterone, control samples were collected at times 0 and 240 min to determine the temporal changes associated with the basal production of progesterone.

**Figure 1.**
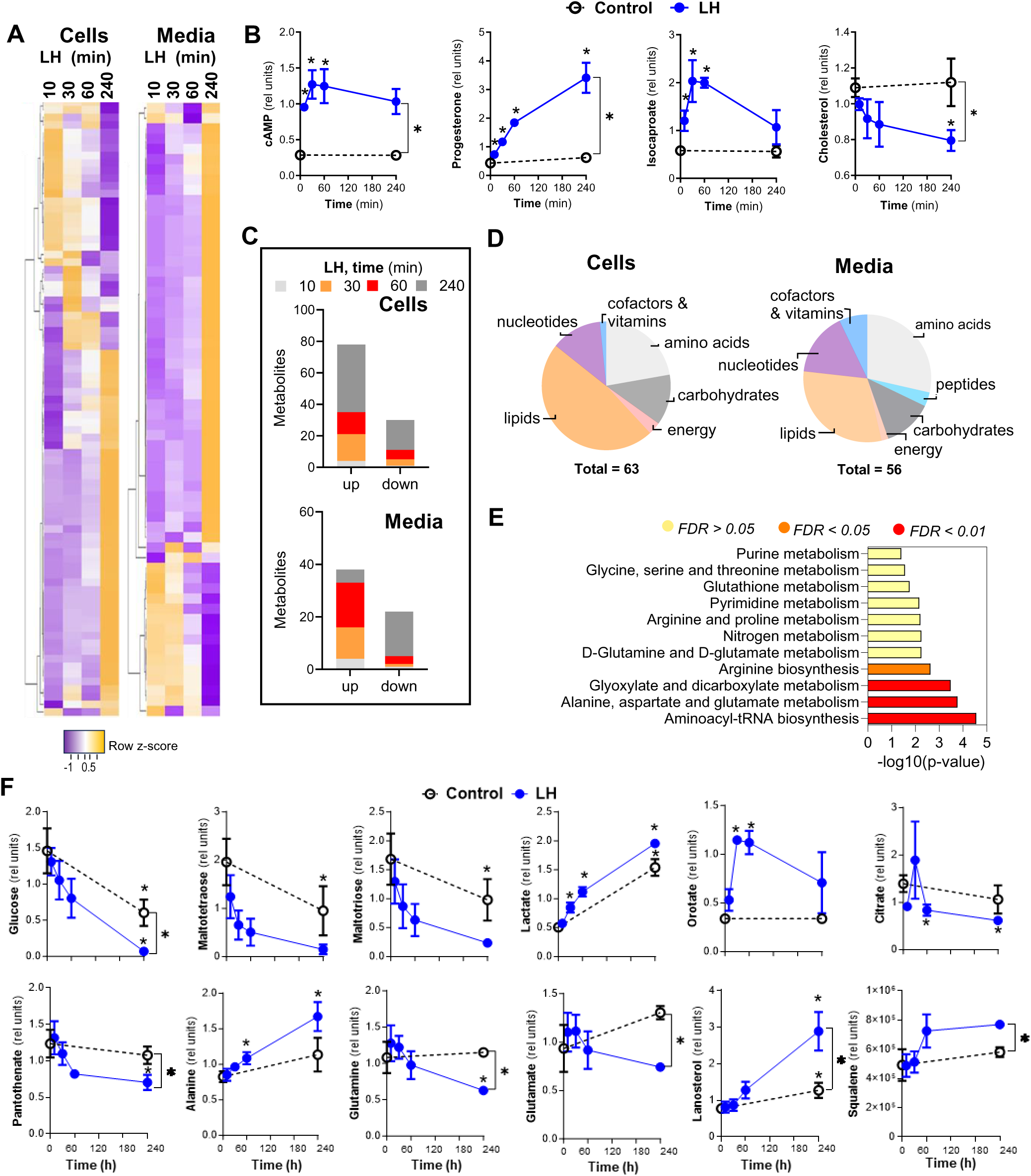
Luteinizing hormone (LH) stimulates acute changes in cellular lipids and glucose metabolism in luteal cells. Enriched preparations of small luteal cells treated with LH (10 ng/ml) for 10-240 min or post-incubation media metabolites were analyzed by mass spectrometry. (**A**) Heatmaps showing the most significant (≥ 1.5 or ≤ −1.5) time-dependent changes post-LH treatment in media and cell extracts. The Heatmap was prepared using http://www.heatmapper.ca/. (**B**) Changes in cellular or media content of cyclic AMP (cAMP), progesterone, isocaproate and cholesterol in small luteal cells treated with LH (10 ng/ml) for 0-, 10-, 30-, 60- and 240-min. Values are presented as relative units (RU) and means ±SEM, n = 3. Asterisk * indicates P ≤ 0.05. (**C-D**) Time-dependent changes in metabolite concentrations and classes in media and cell extracts. (**E**) Metabolic pathways changed post-LH treatment. Analysis was done using MetaboAnalyst (Version 3.0, URL: http://www.metaboanalyst.ca) and the most significantly changed (≥ 1.5 or ≤ −1.5) metabolites in media and cell extracts. False Discovery Rate (FDR) presented with different colors. (**F**) Changes in cellular or media content of selected metabolites in the small luteal cells treated with LH (10 ng/ml) for 0 to 240-min. Values are presented as relative units (RU) and means ±SEM, n = 3. Asterisk * indicates P ≤ 0.05.

Mass spectrometry revealed acute and prominent changes in metabolite concentrations depending on treatment and time in both cell extracts and media (Figure 1A; Supplementary Table 1). As expected, LH rapidly increased cAMP in cells (4.4-fold; P < 0.05) after 10 min of incubation, with maximal levels attained within 30-60 min (Figure 1B). A corresponding rapid and continual increase in progesterone secretion was observed. LH also stimulated an acute elevation (3.5-fold; P < 0.05) in isocaproate, a product of cholesterol cleaved by CYP11A1, the mitochondrial cholesterol side-chain cleavage enzyme, with a concomitant reduction (28% reduction) in total cellular cholesterol content after 240 min (Figure 1B). These findings demonstrate that small steroidogenic luteal cells employed in these studies provide an appropriate model to examine metabolic events activated during the steroidogenic response to LH.

Treatment with LH rapidly altered the luteal cell metabolic profile. Briefly, 276 and 117 metabolites were identified in cell extracts and conditioned culture media, respectively. Incubation with LH for 10 min induced significant changes (P < 0.05) in 5 metabolites in cell extracts and media samples; after 30 and 60 min, the number of metabolites increased to 20, and after 240 min of treatment, it increased to 34. In control samples, 54 metabolites showed significant changes during the 240 min incubation (Figure 1C; Supplementary Table 1). The most significantly changed metabolites in cells and media were categorized as lipids, amino acids, nucleotides, and carbohydrates (Figure 1D). Enrichment analysis indicated significant changes (P < 0.05; FDR < 0.05) in aminoacyl-tRNA biosynthesis; alanine, aspartate, and glutamate metabolism; glyoxylate and dicarboxylate metabolism and arginine biosynthesis (Figure 1E; Supplementary Table 1).

### LH enhances glucose metabolism, TCA cycle and glutamine metabolism

Glucose is a primary energetic substrate in each living cell (29). Once inside the cells, glucose molecules are broken down into pyruvate and lactate. Further, pyruvate can be transported to the mitochondria via mitochondrial pyruvate carriers (MPCs), where it is utilized in the TCA cycle to produce energy and intermediates required for *de novo* lipogenesis or nucleotide synthesis (29).

We observed that LH induced a substantial decrease in the intracellular concentration of glucose (59% reduction, P < 0.05), as well as the glucose storage compounds - maltotriose and maltotetraose (Figure 1F), polysaccharides with glucose molecules linked with α-1,4 glycosidic bonds (86% and 92% reduction, P < 0.05). In media samples, a dramatic drop in fructose and glucose concentrations (90% and 51% reduction, respectively) was found after 30 min with LH treatment (Supplementary Table 1). Simultaneously, a 1.3-fold increase in lactate (P < 0.05) was observed in media samples, evidence of enhanced glycolysis (Figure 1F) (30). Glucose can also be directed to the pentose phosphate pathway (PPP) and hexosamine biosynthetic pathway (HBP), which lead to the synthesis of nucleotides or reduced nicotinamide adenine dinucleotide phosphate (NADPH), and glycolipids, glycoproteins, respectively (31). Herein, LH treatment elevated levels of orotate, a pyrimidine precursor, reaching maximal levels within 30-60 min (3.4-fold increase, P < 0.05) in cells indicating enhanced activity of the PPP in response to LH (Figure 1F). We observed a rapid utilization of citrate (56% reduction, P < 0.05) in cells suggesting increased activity of the TCA cycle (Figure 1F). In addition, at 240 min, LH provoked a depletion (35%) of pantothenate, a precursor of coenzyme A (CoA), which is a cofactor required for acetyl-CoA formation and lipid metabolism (Figure 1F) (32). As citrate can be transported to the cytoplasm and used to produce acetyl-CoA required for lipid synthesis (33), the observed changes suggest enhanced *de novo* lipogenesis. LH raised the concentration of alanine (Figure 1F) and tryptophan (Supplementary Table 1), two amino acids that can be used as a source of pyruvate for the TCA cycle. Beyond pyruvate, the TCA cycle can be replenished by anaplerosis by conversion of glutamine and glutamate to α-ketoglutarate (31). Glutamine and glutamate were significantly reduced (56 and 53%, respectively) in media samples 240 min after LH treatment indicating the possible use of these amino acids to generate TCA intermediates or glutathione, a known antioxidant (34), especially in view of the rapid and persistent decreased levels of cysteine-glutathione-disulfide (CYSH-GSH; 33% reduction, P < 0.05) in cells following LH treatment.

### LH stimulates hydrolysis of phospholipids

Phospholipids which are abundant in cell membranes, can be hydrolyzed by phospholipase A (PLA) to release fatty acids resulting in the formation of lysophospholipids. Fatty acids can be used as substrates for mitochondrial energy production or synthesis of lipid mediators, including prostanoids and leukotrienes or steroids (32). We observed substantial increases in intracellular concentrations of various lysophospholipids at 240 min of LH treatment, suggesting a delayed enhanced phospholipase A2 (PLA2) activity in LH-stimulated cells (Supplementary Table 1). Thus, it is likely that PLA activity is augmented in LH-treated luteal cells, consistent with previous reports of elevated cytosolic PLA2 levels and activity in in granulosa cells of primate periovulatory follicles stimulated with human chorionic gonadotropin (hCG) (35).

### LH rapidly evokes changes in *de novo* cholesterol synthesis

*In vivo*, cholesterol is provided to luteal cells by high- and low-density lipoproteins (HDL and LDL, respectively) (36). Once inside the luteal cell, lipoprotein-derived cholesterol is used for steroidogenesis via an LHCGR/PKA-responsive pool of cholesterol esters (esterified with long-chain fatty acids) stored in lipid droplets (37). Additionally, cholesterol can be synthesized *de novo* in cells from acetyl-CoA produced in mitochondria during the TCA cycle or from acetate (33, 36). Herein, under serum-free culture conditions, lanosterol and squalene, precursors for the *de novo* synthesis of cholesterol, were elevated (2.5- and 1.5-fold, respectively) in cells and media 240 min after LH treatment suggesting enhanced activity of *de novo* lipogenesis and enzymes involved in cholesterol synthesis (Figure 1F).

Our results obtained by mass spectrometry analysis were confirmed by NMR spectroscopy (Supplementary Figure 1A-B), where metabolic changes were analyzed in small luteal cells collected after 240 min of incubation with LH. Herein, compared to controls, we identified augmented concentrations of glycine, alanine, branched-chain amino acids, threonine, and lactate. Simultaneously, we observed depletion in metabolites related with the TCA cycle and redox homeostasis, such as glutamate and glutamine, citrate, aspartate, arginine, glutathione, and choline-containing compounds following LH treatment (Supplementary Figure 1B).

### LH stimulates mitochondrial respiration

Cells produce energy through two major pathways: oxidative phosphorylation (OXPHOS) and glycolysis. During oxidation and reduction reactions, substrates such as glucose, glutamine, and fatty acids are converted within the mitochondria into adenosine triphosphate (ATP) (29, 31). Due to the observed changes in the concentration of glucose and glutamine, we next determined the effect of LH on mitochondrial respiration. The components of the electron transport chain (ETC) in small luteal cells were analyzed using a cocktail of antibodies specific for determining the relative levels of OXPHOS complexes in mitochondria. All five complexes (CI-CV) were detected (Figure 2A). Mitochondrial respiration was measured in real-time as oxygen consumption rate (OCR) in intact primary small luteal cells using the Seahorse Mito Stress Assay during which OCR is measured before (basal respiration) and after injections with inhibitors of the ETC (Figure 2B). Along with elevated concentrations of lactate (1.5-fold increase, P < 0.05) (Figure 1F, 2C), treatment with LH increased basal respiration and ATP production (1.7-, and 2.8-fold, respectively) compared to control cells. No changes were observed in non-mitochondrial respiration (Figure 2D-E). Maximal respiratory capacity measured after FCCP injection was markedly lower (28% reduction) in cells incubated with LH versus control cells, indicating a reduced availability of substrates used for oxidation following treatment with LH (Figure 2E). Furthermore, elevated basal extracellular acidification rate (ECAR) were found in LH treated cells (Supplementary Figure 2A).

**Figure 2.**
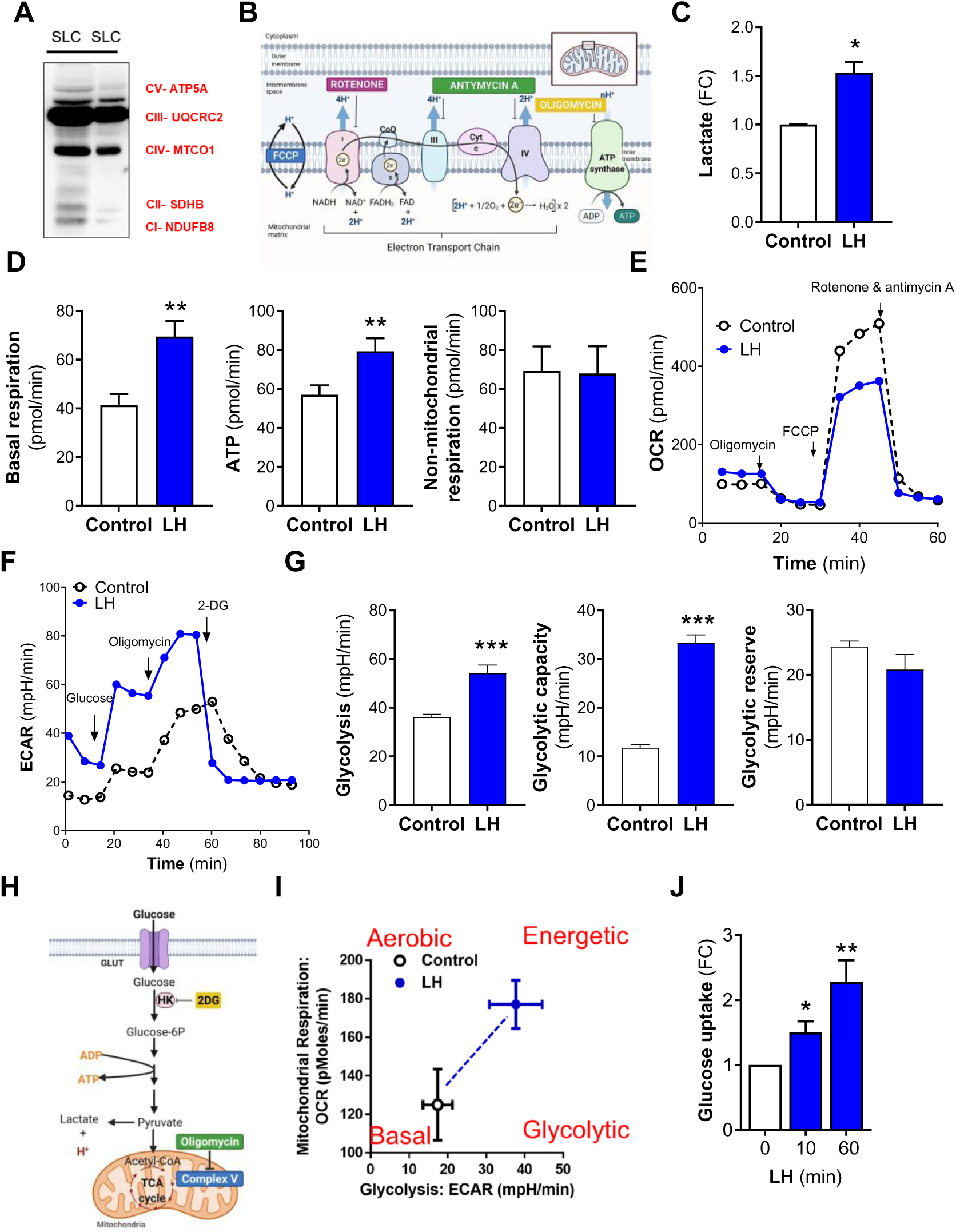
LH affects mitochondrial respiration and glycolysis. (**A**) Representative western blot using a total OXPHOS antibody cocktail showing content of proteins of electron transport chain in the small luteal cells. (**B**) Mitochondrial respiration was analyzed using Seahorse Cell Mito Stress Test. Diagram presenting electron transport chain and function of inhibitors used in Seahorse analysis. Mitochondrial respiration was measured as oxygen consumption rate (OCR) following a sequential addition of inhibitors of mitochondrial function: oligomycin, carbonyl cyanide-ptrifluoromethoxyphenylhydrazone (FCCP), and a combination of rotenone and antimycin A. (**C**) Lactate concentration measured in medium post-Seahorse analysis. Data for control and LH-treated cells are presented as a fold change (FC) and mean ±SEM (n=3). Lactate was measured using ELISA method. Data were analyzed using Student’s t-test. Asterisk * means significant change with P < 0.05. (**D**) Basal respiration, ATP production, and non-mitochondrial respiration in untreated (Control) and LH-treated (60 min) small luteal cells. Data are presented as mean±SEM (n=3). Data were analyzed using Student’s t-test. Asterisks ** indicate P < 0.01. (**E**) Representative graph showing oxygen consumption rate (OCR) in the small luteal cells treated with LH (10 ng/ml) for 60 min (blue solid line) and control cells (black dotted line). (**F**) Representative graph showing changes in ECAR in the small luteal cells untreated (Control) or treated with LH (10 ng/ml) for 60 min. (**G**) Glycolysis, glycolytic capacity, and glycolytic reserve in Control and LH-treated (60 min) cells. Data are represented as mean±SEM (n=3) and were analyzed using Student’s t-test. Asterisks *** indicate significant change with P < 0.001. (**H**) Glycolytic rate was analyzed using Seahorse Glycolytic Rate Assay. Diagram explaining site of action of inhibitors used in Seahorse analysis. Glycolytic rate was measured as extracellular acidification rate (ECAR) following a sequential addition of glucose, oligomycin (inhibitor of ATP synthase) and 2-deoxyglucose (2-DG; inhibitor of hexokinase-HK). (**I**) Bioenergetics phenotype of untreated (Control) and LH-treated cells done basing on OCR and extracellular acidification rate (ECAR) values obtained from Seahorse analysis (n=3). (**J**) Glucose uptake assay done using luminescent method for untreated (Control) and LH-treated cells for 10- and 60-min. Data are presented as a fold change (FC) and mean±SEM (n=3). Data were analyzed using a one-way ANOVA test. Asterisks *, ** mean significant change with P < 0.05 and P < 0.01, respectively.

### LH stimulates glucose uptake and glycolytic capacity

Our mass spectrometry data and previous studies indicate that LH affects glucose uptake and glycolysis (38–41). Thus, we designed experiments to measure the glycolytic rate and glucose uptake in LH-treated small luteal cells. Seahorse analysis using the Glycolysis Stress Assay revealed elevated ECAR in LH-treated cells compared to control cells (Figure 2F). Glucose and oligomycin injection increased ECAR in LH-treated cells versus control, reflecting enhanced glycolysis and glycolytic capacity after gonadotropin stimulus (1.5 and 3-fold, respectively), while the glycolytic reserve remained unchanged (Figures 2F-H). Elevated ECAR and OCR values indicate that LH acutely promotes an energetic phenotype in luteal cells (Figure 2I), which signifies that LH stimulates metabolic pathways that boost the cellular flow of metabolites for both glycolysis and mitochondrial respiration in luteal cells. Utilizing a bioluminescent method based on detecting 2-deoxyglucose-6-phosphate (2DG6P), we observed that LH stimulated significant increases in glucose uptake within 10 min, and uptake continued to rise over 60 min (2.2-fold; Figure 2J). These findings confirm the rapid and stimulatory actions of LH on acute glucose uptake and glycolysis in luteal cells.

Because our previous microarray studies (GSE83524) revealed transcripts for glucose transporters (*GLUT 1*, *3*, *8*, and *13*) in luteal cells, with *GLUT1* being the most highly expressed (Supplementary Figure 2B), we measured glucose uptake in cells pretreated for 60 min with a selective inhibitor of GLUT1 (BAY876; 1 µM) or an inhibitor of GLUT1 and GLUT4 (Fasentin; 100 µM). Treatment with both GLUT inhibitors abolished the stimulatory effect of LH on glucose uptake, indicating the importance of GLUT1 in LH-responsive glucose uptake in luteal cells (Supplementary Figure 2C); however, they did not prevent LH-stimulated progesterone synthesis (Supplementary Figure 2D). Also, inhibition of hexokinase, the enzyme catalyzing the first essential step of glucose metabolism, i.e., conversion of glucose into glucose-6-phosphate (42), using lonidamine (10-100 µM), a selective hexokinase inhibitor, did not affect steroidogenesis in cells treated with LH (Supplementary Figure 2E).

### LH directs glucose through glycolysis, pentose phosphate pathway and hexosamine biosynthetic pathway

We performed a fluxomics analysis to determine how small luteal cells use glucose. Herein, cells were incubated in media containing 5 mM [U^13^C_6_]-labeled glucose and then treated with or without LH (10 ng/ml) for 60 or 240 min. We discerned the time-dependent effects of LH on metabolic changes by using NMR (Figure 3A-B). Incubation with LH rapidly depleted [U^13^C_6_]-glucose from cells (48% reduction) and media (52% reduction) (Figure 3C-D) with the simultaneous increase (by 15-16-fold) in progesterone production (Supplementary Figure 3A; Supplementary Table 2). Bioinformatics analysis revealed the most significant (P < 0.05; FDR < 0.05) enrichment of glucose metabolites in the following pathways: alanine, aspartate, and glutamate metabolism; TCA cycle; glycolysis/gluconeogenesis; galactose metabolism and butanoate metabolism (Figure 3B) suggesting changes within pathways related with production of energy, synthesis of nucleotides and proteins or cofactors catalyzing synthesis of lipids and proper function of steroidogenic machinery (34).

**Figure 3.**
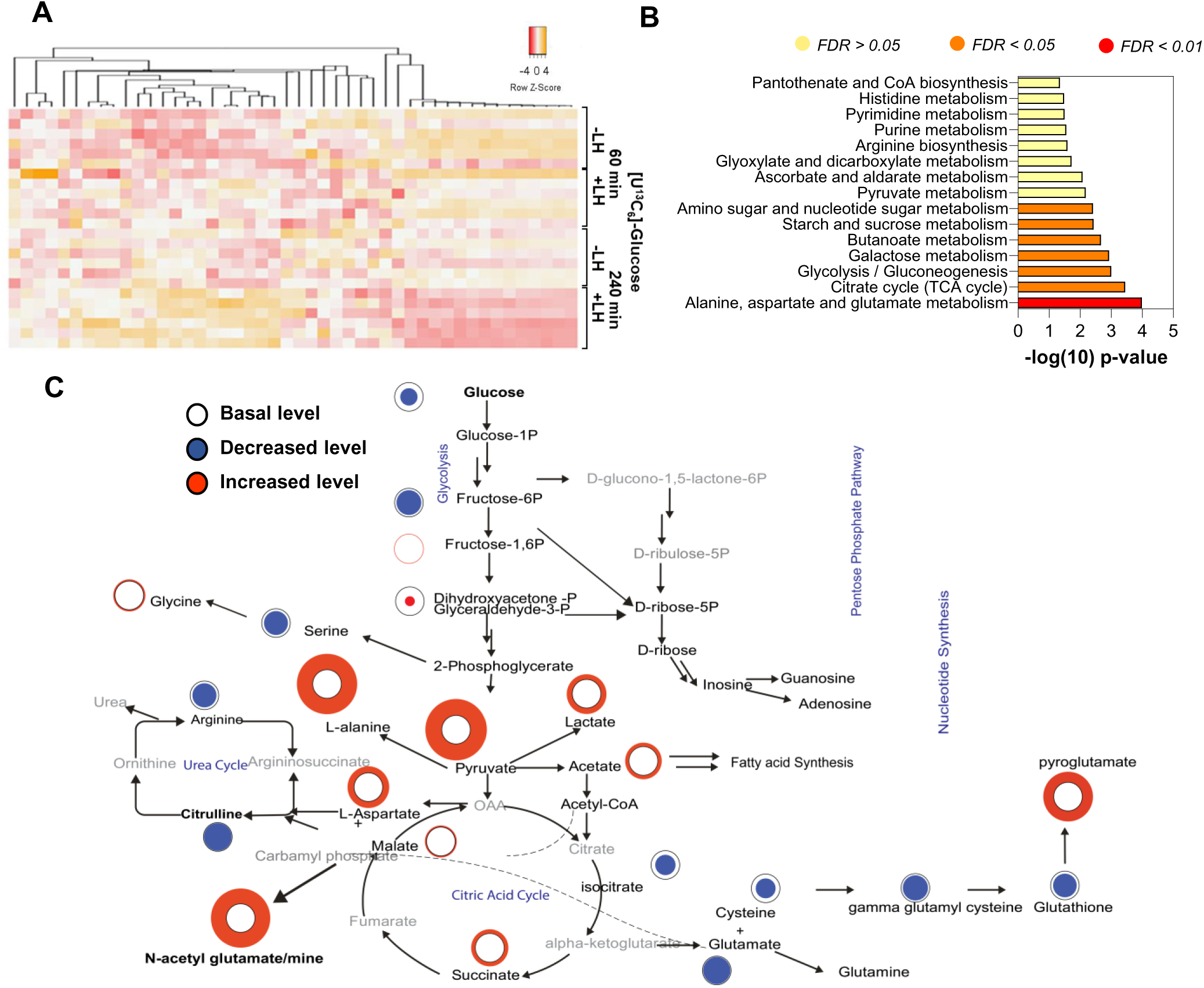
LH stimulates glucose metabolism in the luteal cells. Enriched preparation of small luteal cells treated with [U^13^C_6_]-labeled glucose (5 mM) alone or in the presence of LH (10 ng/ml) for 60- and 240-min. Cellular metabolites were analyzed by Nuclear Magnetic Resonance (NMR) spectroscopy. (**A**) Heatmap representing the most significant changes in [U^13^C_6_]-labeled metabolites. (**B**) Enrichment pathway analysis done using MetaboAnalyst (Version 3.0, URL: http://www.metaboanalyst.ca). Analysis was performed using the most significantly (P < 0.05) changed metabolites in cells and media. (**C**) Flow chart showing the most significant changes and metabolic pathways in cell extracts cultured in the presence of LH. Black, blue, and red circles represent the basic, decreased, or increased concentration of metabolites.

We found a prominent decrease in metabolites related with the glycolytic pathway, such as glyceraldehyde-3-phosphate (38% depletion) and increased production of lactate (1.5-fold increase) (Figure 3C). Incubation with LH also induced acute changes within the PPP and the HBP reflected by elevated concentrations of uridine (3-fold increase), cytidine triphosphate/uridine triphosphate (CTP/UTP) (6.7-fold increase), and uridine diphosphate (UDP)-glucose (4-fold increase) (Figure 3C; Supplementary Figure 3B).

### LH leads glucose flow through TCA cycle, lipid metabolism and redox homeostasis

LH treatment enhanced the concentration of metabolites related with the TCA cycle, such as ^13^C-labeled-pyruvate and -succinate (1.5-fold increase for both), as well as decreased (34% reduction) content of ^13^C-labeled-isocitric acid and malate (Figure 3C; Supplementary Figure 3B). We also observed significantly increased content of nicotinamide adenine dinucleotide (NADH) in LH-treated cells (3.1-fold; Figure 3C; Supplementary Figure 3B), which is produced during glycolysis and the TCA cycle indicating enhanced activity of these pathways (29, 34). There was also an elevated concentration of alanine and aspartate (2.2 and 1.3-fold), two amino acids used in the TCA cycle, gluconeogenesis, or nucleotide synthesis (34). Compared to control, LH decreased the content of carnitine (20% depletion), which transports long-chain fatty acids into mitochondria to be oxidized for ATP production, while increasing the content of CoA (1.3-fold change) and acetate (1.3-fold increase) (Figure 3C; Supplementary Figure 3B), suggesting enhanced activity in pathways supplying cells with acetyl-CoA. There were also changes in metabolites related to redox homeostasis, including an increase in glutathione (2.0-fold increase) concentration (Figure 3C; Supplementary Figure 3B). A summary of the flux of [U^13^C_6_]-labeled glucose in response to LH is shown in Figure 3D.

### Inhibition of pyruvate synthesis and TCA cycle activity abrogates LH-stimulated steroidogenesis

According to our previous microarray data (GSE83524) (18, 27), there is a prominent increase in the expression of genes encoding factors related with cholesterol (*APOA1*, *APOD*, *APOE*, *HMGCR*, *HMGCS1*, *SQLE, ACAT2*) and fatty acids synthesis or metabolism (*ACLY*, *ACSS2*, *FASN, CRAT, ACSL3, ACSL4*) in small and large luteal cells in comparison to theca and granulosa cells (Figure 4A). We also observed elevated expression of genes encoding glucose transporters (Supplementary Figure 2B) and glucose metabolism (*ALDOA*, *LDHB*, *G6PD*, *PGD, IDH1, IDH3G*) during the follicle-to-luteal transition (Figure 4A).

**Figure 4.**
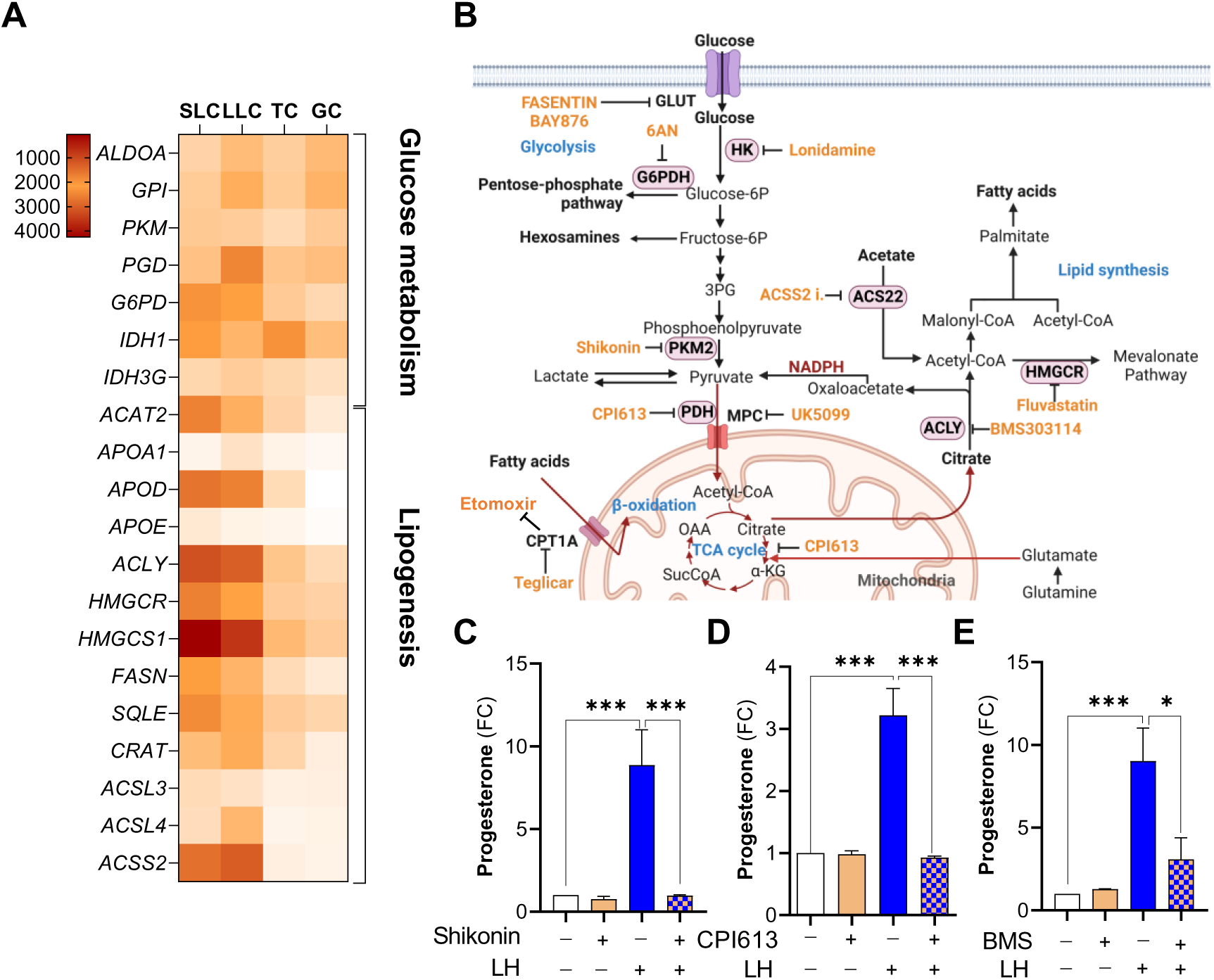
Glycolysis, TCA cycle and *de novo* synthesis of fatty acids are vital for steroidogenic capacity of small luteal cells. Small luteal cells were pretreated with inhibitors of selected metabolic pathways for 60 min and then treated with LH (10 ng/ml) for 240 min. Progesterone concentration was measured in media samples. (**A**) Heatmap representing the expression of genes related to lipid metabolism in granulosa and theca cells (GC and TC) as well as small and large luteal cells (SLC and LLC). (**B**) Flow chart showing sequence of metabolic events occurring in cells with information of inhibited enzymes and applied inhibitors. (**C-E**) Progesterone production by small luteal cells pretreated with inhibitors [PKM2-Shikonin (1 μM), PDH and KDGH-CPI613 (25 μM) or ACLY-BMS30314 (25 μm)]. Data are presented as a fold change (FC) and mean ± SEM (n=3-4). Data were analyzed using one way ANOVA test. Asterisks *, *** mean significant change with P < 0.05, P < 0.001.

Further, we conducted experiments to determine the potential role of metabolic pathways related to glucose and lipid metabolism on the steroidogenic response to LH (Figure 4B). Because we observed LH-stimulated effects of glucose flow through the PPP, we treated luteal cells with 6-aminonicotinamide (6-AN) to block glucose-6-phosphate dehydrogenase (G6PD), an enzyme necessary for directing glucose to the PPP and responsible for NADPH production (34). Pretreatment with this inhibitor did not block LH-simulated progesterone synthesis despite applying relatively high concentrations of the compound (10-500 µM) (Supplementary Figure 4A). Next, we inhibited pyruvate kinase 2 (PKM2), catalyzing the last step in glycolysis, i.e., dephosphorylation of phosphoenolpyruvate to generate pyruvate (Figure 4B). Pretreatment with increasing concentrations (0.1-5 µM) of the PKM2 inhibitor-Shikonin decreased LH-stimulated progesterone production (Figure 4C; Supplementary Figure 4B). Treatment with Shikonin also reduced LH-stimulated ECAR, glycolysis, glycolytic capacity, and glycolytic reserve (Supplementary Figure 4C-D). Since luteal cells treated with LH showed prominent changes within the TCA cycle, we also employed CPI-613, an inhibitor of pyruvate dehydrogenase (PDH) and α-ketoglutarate dehydrogenase (α-KGDH) (43) catalyzing conversion of pyruvate to acetyl-CoA and conversion of α-ketoglutarate to succinyl-CoA, respectively (Figure 4B). We observed that pretreatment with CPI-613 (25 µM) profoundly decreased progesterone production in cells treated with LH (Figure 4D). These results suggest a crucial role of glycolysis and pyruvate production, as well as the TCA cycle in luteal steroidogenesis.

### Fatty acids are important source of energy and fundamental for LHCGR/PKA signaling and steroidogenesis in luteal cells

Our global metabolomics analysis identified a rapid depletion of citrate in luteal cells treated with LH. Citrate produced in TCA cycle can be transported to the cytoplasm and cleaved to acetyl-CoA and oxaloacetate by ATP citrate lyase (ACLY) with concomitant hydrolysis of ATP to adenosine diphosphate (ADP) and phosphate. The generated oxaloacetate can be converted to malate and pyruvate with NADPH production catalyzed by malate enzyme. Acetyl-CoA can also be produced from acetate in the reaction catalyzed by acetyl-CoA synthetase short-chain family member 2 (ACSS2) (44). The product, acetyl-CoA, is a substrate for fatty acid and cholesterol synthesis (Figure 4B). Genes encoding both ACLY and ACSS2 are abundant in luteal cells (Figure 4A). Pretreatment for 60 min with a specific ACSS2 inhibitor (10 µM) prior to treating luteal cells with LH for 240 min did not change LH-stimulated progesterone production (Supplementary Figure 5A). To determine whether cholesterol synthesis supports the acute steroidogenic response to LH, we inhibited 3-hydroxy-3-methylglutaryl-CoA reductase (HMGCR), an enzyme considered the rate-limiting enzyme for this pathway (Figure 4B). Incubation of luteal cells in the presence of fluvastatin (1-50 µM), a clinically used inhibitor of HMGCR (45), did not affect LH-stimulated progesterone production, suggesting that fatty acid synthesis, instead of cholesterol synthesis, may have an essential role for the maintenance of steroidogenic capacity of luteal cells (Supplementary Figure 5B). Thus, we focused our further experiments on the function of ACLY.

Pretreatment with increasing concentrations of ACLY inhibitor, BMS303141 (BMS, 5-50 µM), caused concentration-dependent reductions in the stimulatory effect of LH on progesterone production in small luteal cells (20-65% of reduction), implicating *de novo* lipogenesis in the steroidogenic response (Figure 4E; Supplementary Figure 5C). We next determined whether mitochondria utilize endogenous fatty acids to generate ATP and if fatty acids are required for progesterone production in luteal cells. We treated cells with etomoxir (30 µM), an inhibitor of carnitine palmitoyltransferase 1A (CPT1A), which transports fatty acids into mitochondria (32) (Figure 4B) and measured OCR using Seahorse (Figure 5A). Pretreatment with etomoxir significantly inhibited the stimulatory effects of LH on OCR (Figure 5A), ATP production (Figure 5B, 80% reduction) and progesterone synthesis (Figure 5C), implicating fatty acids as important mediators of luteal function (Figure 5A-C; Supplementary Figures 5D). This finding was confirmed by pretreatment with another inhibitor of CPT1A, Teglicar, where increasing concentrations of Teglicar (5-50 μM) decreased LH-stimulated progesterone (10-57% reduction), and ATP production (42-76% reduction) measured using luminescence (Figures 5C-D). Furthermore, genetic knockdown of *CPT1A* also inhibited (P < 0.01) LH-stimulated progesterone production (53% of reduction) (Figure 5E). Knockdown of CPT1A did not change the content of mitochondrial OXPHOS proteins and TOM20 or steroidogenic machinery proteins (STAR, CYP11A1) (Figure 5F).

**Figure 5.**
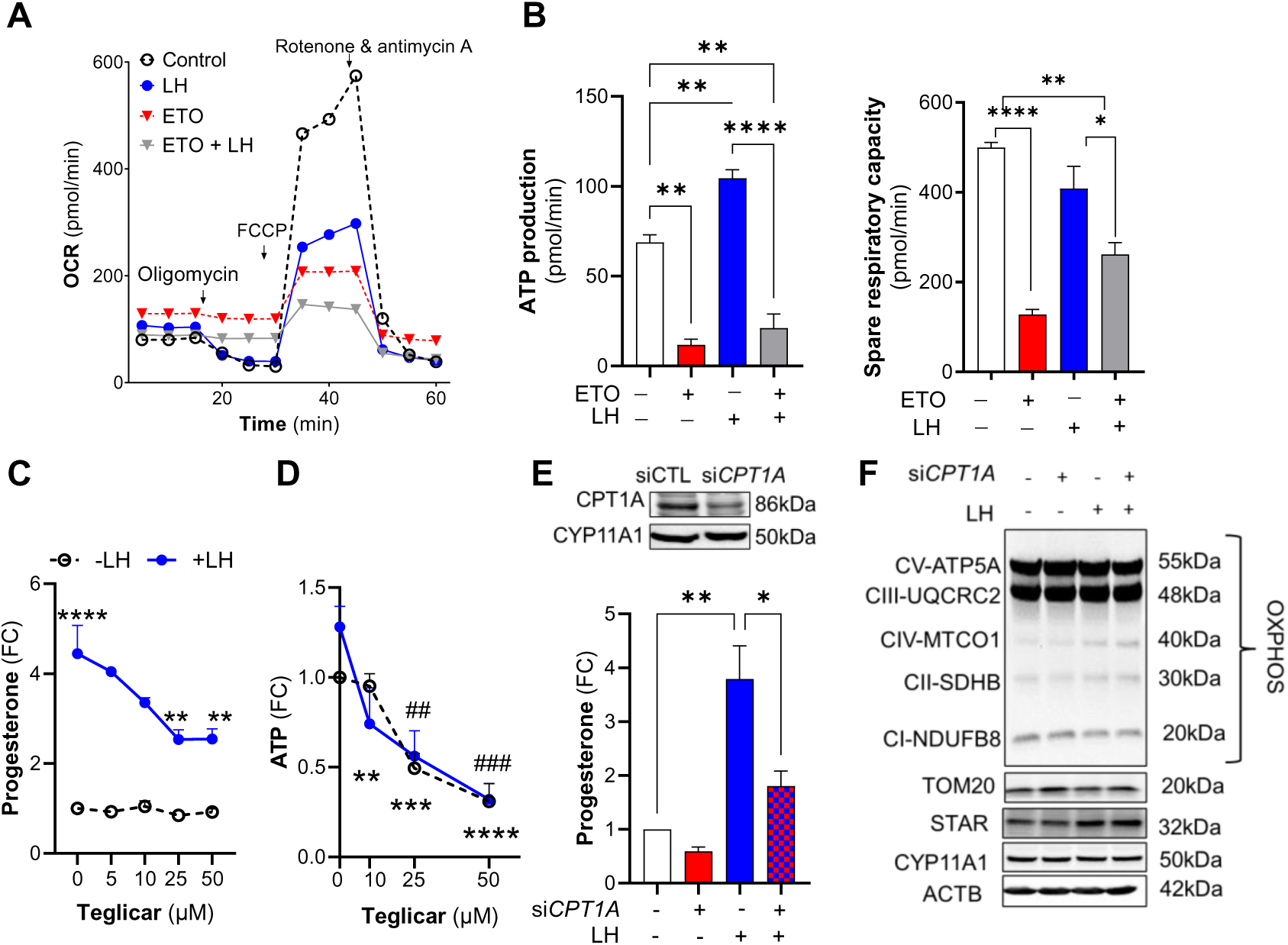
Endogenous fatty acids are an essential source of energy in luteal cells. (**A**) Representative graph showing oxygen consumption rate (OCR) in the small luteal cells pretreated with etomoxir (ETO; 30 μM) and then treated with LH (10 ng/ml) for 60 min. (**B**) ATP production and spare respiratory capacity in the small luteal cells pretreated with etomoxir (ETO) and then treated with LH. Data are represented as mean±SEM (n=6-8) and were analyzed using one-way ANOVA test. Asterisk *, ** and **** indicates P < 0.05, P < 0.01, and P < 0.0001, respectively. (**C**) Dose-dependent effects of CPT1A inhibitor (Teglicar; 5-100 μM) on progesterone production by untreated (Control) and LH-treated small luteal cells. Data are represented as fold change (FC) and mean±SEM (n=3-5). Data were analyzed by using a two-way ANOVA. Asterisks **, ***, **** represent significant change with P < 0.05, P < 0.01, P < 0.0001, respectively. (**D**) ATP production by luteal cells pretreated with CPT1A inhibitor (Teglicar; 10-50 μM) and then treated with LH (10 ng/ml) for 240 min. ATP production was measured by using luminescence. Data are presented as fold change and mean±SEM (n=3-5) and were analyzed by using two-way ANOVA test. Asterisks **, ***, **** mean significant change with P < 0.05, P < 0.01, and P < 0.001, respectively for cells treated with LH alone *versus* Teglicar+LH. Symbols ^##^ and ^###^ represent significant change with P < 0.01 and P < 0.01, respectively for cells treated with Teglicar *versus* Control (untreated). (**E**) Progesterone production by small luteal cells with knockdown CPT1A (si*CPT1A;* 50 nM) and treated with LH (10 ng/ml) for 240 min. Control cells were transfected with siControl (50 nM). Data are presented as a fold change (FC) and mean±SEM (n=3-5). Data were analyzed by using a one-way ANOVA test. Asterisks * and ** mean significant change with P < 0.05 and P < 0.01, respectively. Efficiency of siRNA transfection was confirmed using western blotting and is presented above the bar graph. (**F**) Representative blots showing content of electron transport chain (ETC) proteins in the small luteal cells with knockdown CPT1A (si*CPT1A*; 50 nM) and then treated with LH (10 ng/ml) for 240 min.

### LHCGR/PKA pathway triggers phosphorylation of enzymes involved in *de novo* lipogenesis

Considering crucial role of fatty acids in LH-mediated stimulatory effects on progesterone by highly steroidogenic luteal cells, we tested effects of LH on phosphorylation of enzymes involved in fatty acids synthesis. PKA regulates the phosphorylation and activation state of at least two enzymes involved in *de novo* lipogenesis - acetyl-CoA carboxylase (ACACA) and ACLY (32, 46). To determine whether LH and PKA triggers the phosphorylation of ACACA or ACLY in steroidogenic luteal cells, we treated cells with increasing concentrations of LH (1-10 ng/ml) or forskolin (FSK; 10 μM), an activator of adenylyl cyclase, for up to 30 min. LH and FSK significantly stimulated the phosphorylation of ACLY at Ser455, a site specific for enzyme activation (47). Phosphorylation of ACLY was enhanced 3-fold at the lowest concentration of LH (1 ng/ml) and the extent of phosphorylation continued to rise with higher concentrations of LH reaching the maximal level (5.4-fold increase) after treatment with 10 ng/ml LH (Figure 6A-B; Supplementary Figure 4F). Both LH and FSK decreased (30-74% of reduction) phosphorylation of ACACA at Ser79, a site specific for enzyme inactivation (48) via inhibition of AMPK (21, 49). Treatment with a PKA inhibitor, H89 (10 µM), prevented LH mediated effects on phosphorylation of ACLY and ACACA in luteal cells confirming that phosphorylation sites are mediated by LH/PKA signaling in luteal cells (Supplementary Figure 5F).

**Figure 6.**
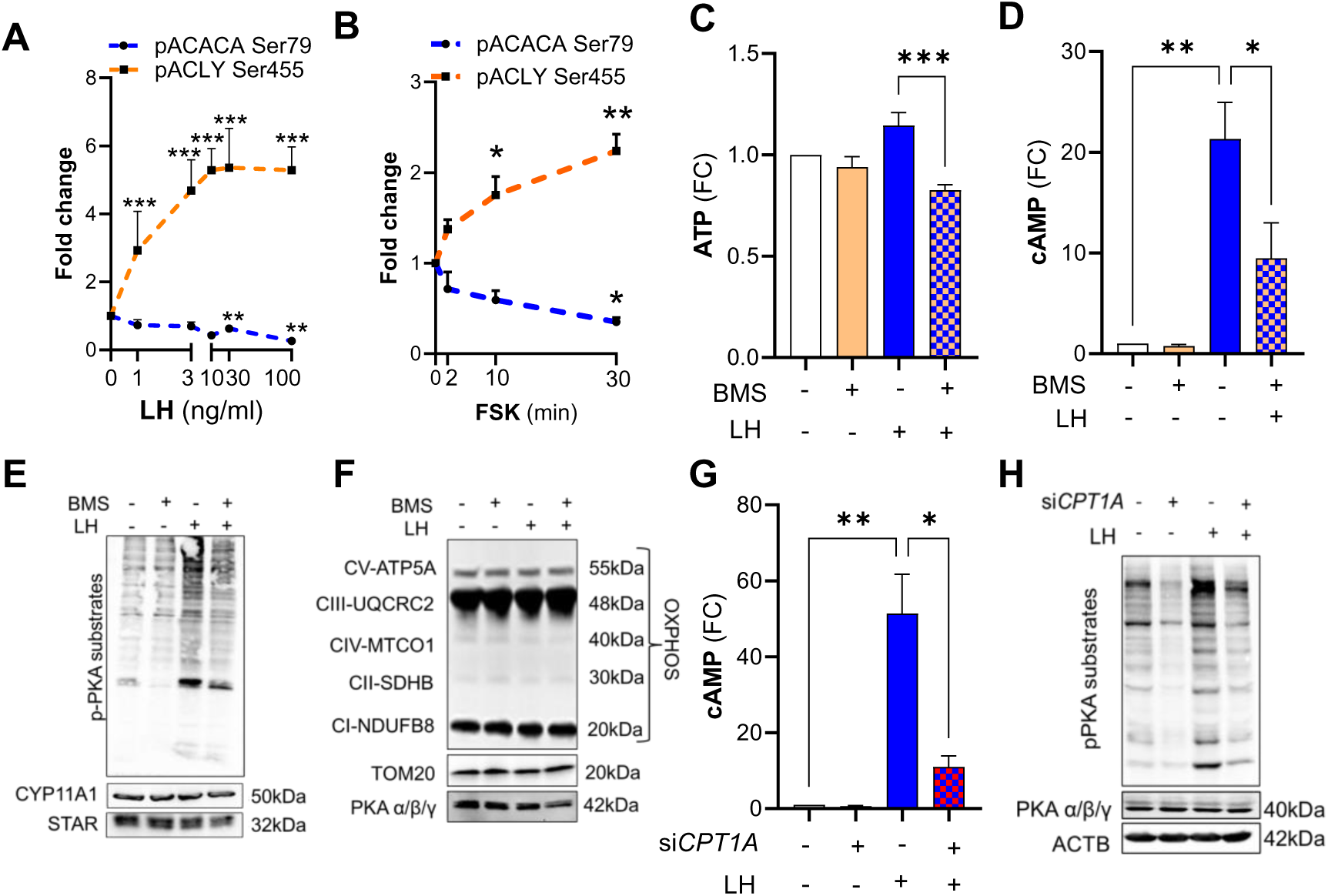
LHCGR/PKA pathway triggers phosphorylation of enzymes involved in *de novo* lipogenesis. (**A**) Phosphorylation of ACC1 Ser79 and ACLY Ser455 in the small luteal cells treated with different concentrations of LH (1-100 ng/ml) for 30 min. Data are presented as fold change and mean±SEM (n=3-5). Data were analyzed using a one-way ANOVA test. Asterisks ** and *** mean significant change with P < 0.01 and P < 0.001, respectively. (**B**) Phosphorylation of ACC1 Ser79 and ACLY Ser455 in the small luteal cells treated with cAMP/PKA activator-forskolin (FSK; 10 μM) for 2-30 min. Data are presented as a fold change (FC) and mean±SEM (n=3-4). Asterisks * and ** mean significant change with P < 0.05 and P < 0.01, respectively. (**C**) ATP production measured by luminescence method in the small luteal cells pretreated with ACLY inhibitor (BMS303141; 10-50 µm) and then treated with LH (10 ng/ml) for 240 min. Data presented as a fold change (FC) and mean±SEM (n=3-5). Data were analyzed by using a two-way ANOVA test. Asterisks **, *** and **** mean significant change with P < 0.05, P < 0.01 and P < 0.001, respectively. (**D**) cyclic AMP (cAMP) production by the small luteal cells pretreated with ACLY inhibitor (BMS303141; 25 µm) and then treated with LH (10 ng/ml) for 240 min. Data presented as fold change (FC) and mean±SEM (n=3-5). Data were analyzed by using a one-way ANOVA test. Asterisks * and **mean significant change with P < 0.05 and P < 0.01, respectively. (**E**-**F**) Representative blots showing phosphorylation of PKA substrates and content of steroidogenic proteins (STAR, CYP11A1), electron transport chain proteins or marker of mitochondria (TOM20) and PKA catalytic subunits in the small luteal cells pretreated with ACLY inhibitor (BMS303141; 25 µM) and then treated with LH (10 ng/ml) for 240 min. (**G**) cyclic AMP (cAMP) production by the small luteal cells with knockdown CPT1A (si*CPT1A*; 50 nM) and then treated with LH (10 ng/ml) for 240 min. Data are presented as fold change (FC) and mean±SEM (n=3-5). Data were analyzed by using one-way ANOVA. Asterisks ** and *** mean significant change with P < 0.05 and P < 0.01, respectively. (**H**) Representative blots showing phosphorylation of PKA substrates and content of PKA catalytic subunits in the small luteal cells with knockdown CPT1A (si*CPT1A*; 50 nM) and then treated with LH (10 ng/ml) for 240 min.

**Figure 7.**
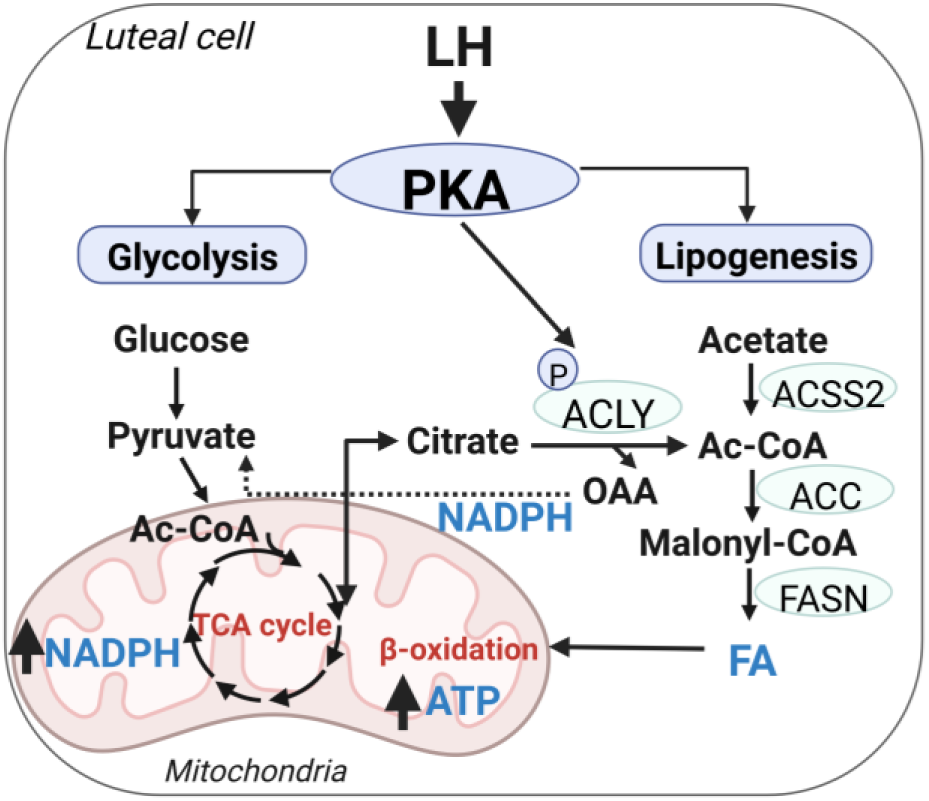
Central Role for Glycolysis and Fatty Acids in LH-responsive Progesterone Synthesis. LH via protein kinase A (PKA) activates metabolic pathways leading to production of acetyl-CoA (Ac-CoA) and *de novo* fatty acid (FA) synthesis. AC-CoA can be obtained from citrate or acetate via ACLY or ASCC2, respectively. FA can be used for β-oxidation and citrate can be used to produce pyruvate with simultaneous production of NADPH.

Next, we tested if ACLY mediates an important function in energy production by luteal cells. Blockage of ACLY reduced basal and LH stimulated ATP and cAMP production (30% and 60% reduction) as well as phosphorylation of PKA substrates without changes in the content of steroidogenic machinery proteins such as CYP11A1 or steroidogenic acute regulatory protein (STAR) as well as OXPHOS and mitochondrial protein (TOM20) or catalytic subunits of PKA α/β/γ (Figures 6C-F; Supplementary Figure 6A). Knockdown or inhibition of CPT1A also abrogated LH-stimulated cAMP production (77% of reduction) and phosphorylation of PKA substrates without changes in the content of catalytic subunits of PKA α/β/γ (Figure 6G-H; Supplementary Figure 6B-C). Similarly, pretreatment with ACLY or CPT1A inhibitor abrogated stimulatory effects of PKA activator, forskolin (FSK; 10 µM) on cAMP, phosphorylation of PKA substrates, and progesterone production without changes in content of STAR (Supplementary Figure 6D-G).

## 3. Discussion

LH-driven alterations in the metabolism of luteal tissue have been recognized for more than 50 years (38, 39). However, these early studies referred only to changes in selected metabolites, such as glucose or lipids in the whole luteal tissue, without providing information on global metabolic changes induced by LH in highly steroidogenic luteal cells (39, 49, 50). In the present study, by using complementary unbiased metabolic approaches, we identified prominent metabolic pathways acutely affected by LH in purified populations of highly steroidogenic primary small luteal cells. We also analyzed the flux of [U-^13^C_6_]-glucose and energetic pathways in LH-stimulated luteal cells. The use of inhibitors of selected metabolic routes allowed the determination of the relative importance of specific metabolic pathways such as glycolysis, TCA cycle, and lipogenesis in the acute response to LH, a vital hormone responsible for the formation, development, and maintenance of luteal function and proper steroidogenesis (51–53). Herein we also provide evidence for LH-mediated post-translational modifications of enzymes involved in *de novo* lipogenesis and the role of *de novo* fatty acid synthesis and fatty acid oxidation in energy homeostasis and LHCGR/PKA signaling and consequently progesterone production.

The findings in this study indicate that LH acutely enhances glucose uptake and glycolysis in the steroidogenic small luteal cells. Previous studies showed that gonadotropins stimulate glycolysis in the primary granulosa cells of sheep, bovine, macaque, human or mice (40, 41, 54–60) as well as mice or rats ovarian follicles cultured or grown in vitro (41, 61) or luteal tissue. In contrast to previous studies in the cow (62), inhibition of glucose uptake and blockage of hexokinase, an enzyme catalyzing conversion of glucose to glucose-6-phosphate, did not affect progesterone production by luteal cells. Since cellular metabolism is an adaptable and flexible network, inhibition of these steps of glycolysis could activate mechanisms overcoming effects of enzyme inhibition. For example, a decrease in the supply of exogenous glucose could enhance the hydrolysis of storage carbohydrates such as maltotriose and maltotetraose or fructose, such as observed in this study, but this remains to be experimentally tested. Caution is needed with this speculation because *in vivo* treatment with 2-deoxy-D-glucose, an inhibitor of hexokinase, prevented both the occurrence of the estrous cycle and the formation of the corpus luteum in cows (63). In contrast, blockage of PKM2, an enzyme that catalyzes the final step of aerobic glycolysis, *i.e.,* dephosphorylation of phosphoenolpyruvate to pyruvate with the simultaneous generation of ATP (29), abrogated LH-mediated stimulatory effects on steroidogenesis in luteal cells. Pyruvate can also be obtained from the sialic acid pathway and catabolism of glutamine, alanine, cysteine, serine, tryptophan, and threonine. As we observed an elevation of alanine and threonine in response to LH, we speculate these amino acids are not the main substrates to fuel the TCA cycle in luteal cells but may be instead used for anabolic processes (34, 64). Notably, inhibition of PKM2 reduced glycolysis confirming effective action of this inhibitor. Glycolysis supports cellular levels of NAD and NADH, cofactors required for proper oxidation occurring in mitochondria (64); therefore, we posit that LH enhances aerobic glycolysis that supplies cells with cofactors and pyruvate required for the proper function of the TCA cycle and mitochondrial oxidation.

Our untargeted metabolomic and fluxomic analysis revealed increases in the concentrations of pyrimidines and purines in LH-treated cells. We also noticed changes within HBP, a branch of glycolysis related to producing UDP-glucose, an essential substrate for protein or lipid glycosylation. LH stimulated the activity of G6PD in rat or bovine luteal explants, and therefore G6PD was identified as the most effective dehydrogenase in the luteal production of NADPH, a cofactor required for progesterone synthesis (65, 66). We did not measure enzyme activity; however, experimental conditions that blocked G6PD did not affect progesterone production, despite high concentrations of inhibitor applied in the experiments. Thus, it seems reasonable to think that the PPP is not required for the regulation of basal or acute luteal progesterone production in response to LH and may rather serve a crucial role in the translation or synthesis of signaling molecules such as glycolipids, proteoglycans, or glycoproteins (31, 67, 68).

The mitochondrial TCA cycle yields cofactors fundamental for the proper function of the ETC, which is required for ATP production. The TCA cycle also provides precursors for various biosynthetic pathways, such as lipid or nucleotide synthesis, and supplies cells with NADPH (68). Inhibition of succinate dehydrogenase activity, an enzyme of the TCA cycle, stimulated G6PD activity, and simultaneously led to the accumulation of cholesterol in the immature rat ovary (66). Because cholesterol is cleaved to pregnenolone by CYP11A1, which uses NADPH as a cofactor, the results of previous studies indicated ineffective NADPH output and impaired activity of CYP11A1 when the TCA cycle was interrupted. In the present study, untargeted metabolomics and fluxomics analysis with labeled [U-^13^C_6_]-glucose revealed significant LH-induced changes in TCA cycle metabolites. Pretreating luteal cells with CPI613, an inhibitor of pyruvate dehydrogenase (PDH) and α-ketoglutarate dehydrogenase (α-KGDH), blocked LH-stimulated progesterone synthesis, revealing a crucial role for the TCA cycle in steroidogenesis. Based on our results, we speculated that the TCA cycle constitutes a leading source of NADPH required for CYP11A1 activity and cofactors essential for maintaining the proper function of ETC in luteal cells. Notably, an intact ETC is crucial to sustain luteal steroidogenesis as inhibiting the ETC complexes also abrogated the LH-stimulatory effects on progesterone production by luteal cells.

Citrate produced within the TCA cycle can be transported to the cytoplasm and cleaved by ACLY to form acetyl-CoA and oxaloacetate. The product, acetyl-CoA, can be used to synthesize fatty acids or cholesterol as well as post-translational modifications of proteins (26, 67). Acetyl-CoA can also be obtained from acetate by ACSS2 (31). Herein, LH rapidly depleted citrate and stimulated the incorporation of labeled-glucose carbons into acetate. Incorporation of [U-C^14^]-labeled acetate into intra-luteal fatty acids and progesterone was observed previously in rat luteal tissue following injection with LH (43). Interestingly, LH and Forskolin, a cAMP/PKA activator, targeted the phosphorylation of ACLY and ACACA at sites required for enzymes activity. ACLY and ACACA are enzymes catalyzing the first step of lipogenesis and fatty acids synthesis (69, 70). In the present study, blockage of ACSS2 did not affect LH-promoted progesterone production by luteal cells. However, inhibition of ACLY inhibited the stimulatory effects of LH on progesterone production, as well as the production of ATP and cAMP, and the phosphorylation of PKA substrates in luteal cells. These results emphasize the importance of ACLY in LH-stimulated energy production, which is essential for the proper function of LHCGR/PKA signaling and steroidogenic capacity of luteal cells.

Our findings show LH-mediated utilization of pantothenate, required for CoA formation, and incorporation of [U-^13^C_6_]-glucose-derived carbons into CoA, a cofactor essential for acetyl-CoA and lipid synthesis (32). Furthermore, LH stimulated the incorporation of [U-^13^C_6_]-glucose-derived carbons into carnitine. Collectively, these findings suggest enhanced production and utilization of fatty acids synthesized from glucose. Fatty acids constitute an essential energy source and substrate for synthesizing different lipid mediators, including prostaglandins, leukotrienes, and steroid hormones (32). Fatty acids are transported to the mitochondria via the mitochondrial carnitine system, where they are oxidized. This transport system consists of the malonyl-CoA sensitive carnitine palmitoyltransferase I (CPT-I) localized in the mitochondrial outer membrane, the carnitine:acylcarnitine translocase, an integral inner membrane protein, and carnitine palmitoyltransferase II localized on the matrix side of the inner membrane (32). CPT1A is responsible for the formation of acylcarnitines by catalyzing the transfer of the acyl group of a long-chain fatty acyl-CoA from CoA to l-carnitine. The product is often palmitoylcarnitine, but other fatty acids may also be substrates (71). To determine whether fatty acids are utilized for energy and steroid synthesis, we interrupted CPT1 activity. Either inhibition of CPT1A by using small molecule inhibitors, or knockdown of *CPT1A* significantly blocked LH-stimulated progesterone production without changes in the content of steroidogenic machinery and mitochondrial proteins. Concurrent with inhibition of CPT1 were decreases in the LH-promoted production of ATP and cAMP as well as phosphorylation of PKA substrates. Thus, we propose that endogenous fatty acids are required for sufficient mitochondrial energy production to fuel LHCGR/PKA signaling in luteal cells.

In the present study, we documented luteal cholesterol utilization reflected by a rapid increase in the concentration of isocaproate, the sidechain of cholesterol cleaved by the mitochondrial CYP11A1 enzyme, and the acute and sustained increase in progesterone production. The conditions for metabolic analysis precluded incubations with serum (which contains lipoproteins), thus removing extracellular cholesterol as a source of cholesterol for progesterone synthesis. We also observed a slight but significant decrease (28%) in total cellular cholesterol content after 240 min of incubation with LH. Those changes were accompanied by elevated intraluteal content of cholesterol precursors, i.e., lanosterol and squalene, suggesting the induction of *de novo* cholesterol synthesis. Activation of *de novo* cholesterol synthesis has recently been reported in mouse ovaries during the LH surge and blocking of SREBP disrupted steroid synthesis (72). In contrast, in fully differentiated luteal cells blocking HMGCR activity, a crucial enzyme in cholesterol synthesis, did not affect LH-stimulated progesterone production. *De novo* cholesterol synthesis is an anabolic process, and although the enzymes involved in the cholesterol synthesis are abundant in the mature corpus luteum (73), *de novo* cholesterol synthesis does not appear to be rate limiting for progesterone synthesis in small luteal cells. Our findings support the notion that this pathway is not a leading source of cholesterol for steroidogenesis in luteal cells (36). This is further supported by our recent findings showing that in response to LH, a pool of cholesterol esters stored in lipid droplets of bovine small luteal cells (21, 22) is hydrolyzed by PKA-activated hormone-sensitive lipase (HSL, also known as LIPE). The released cholesterol is trafficked to the mitochondria for progesterone synthesis. Of note, inhibition of HSL-abrogated lipoprotein stimulated progesterone synthesis in the presence and absence of LH, indicating that the stored cholesterol esters are the hormone-sensitive reservoir of cholesterol and crucial for LH-stimulated progesterone synthesis (22).

The results of this study show for the first time that LH, the main luteotrophic hormone in domestic animals and humans, evokes acute global metabolic changes which support progesterone production by luteal cells. Considering that LH stimulated changes in various branches of glucose metabolism such as glycolysis and the TCA cycle as well as induced post-translational modifications of ACLY and ACACA, enzymes involved in *de novo* lipogenesis, it seems possible that LH induces synthesis of *de novo* fatty acid, which is a primary source of energy, required for proper LHCGR/PKA signaling, and consequently progesterone production. Since the LHCGR signaling pathway is conserved among different animal species and humans, these findings may have a translational impact.

The study evaluated the acute response to LH (up to 4 hr) making additional studies necessary to determine the longer tropic responses to LH. A limitation of the current study is the employment of in vitro incubations of primary cells under serum-free conditions.

## 4. Material and methods

### Reagents

Bovine LH was purchased from Tucker Endocrine Research Institute (Atlanta, GA). Protease and phosphatase inhibitor cocktails were purchased from Sigma Chemical Co. (St. Louis, MO). M199 and fetal bovine serum (FBS) were obtained from Cambrex (Walkersville, MD). Type II collagenase was obtained from Atlantic Biologicals (Lawrenceville, GA). Forskolin were from EMD Chemicals, Inc. (Gibbstown, NJ). The Bio-Rad protein assay kit was purchased from Bio-Rad Laboratory (Richmond, CA). Mito Stress and Glycolysis kit were purchased from Agilent Technologies, Inc. (Santa Clara, CA). H89, UK5099, Shikonin, CPI613, BMS303141, 6AN, Fluvastatin, Lonidamine were purchased from Tocris Bioscience (Minneapolis, MN). ACSS2 inhibitor were obtained from Selleck Chemicals LLC (Houston, TX). Etomoxir (sodium salt) was obtained from Cayman Chemical (Ann Arbor, MI). Progesterone assay was purchased from Diagnostic Systems Laboratories Inc. (Webster, TX). [U-^13^C_6_]-glucose was purchased from Sigma Chemical Co. (St. Louis, MO). Glucose uptake assay was obtained from Promega (Madison, WI). phospho-ACLY (Ser455; #4331), phospho-ACACA (Ser79; #3661), CYP11A1 (#14217), phospho-PKA substrates (#9624), TOM20 (#42406) antibodies were purchased from Cell Signaling Technology, Inc. (Danvers, MA). Total OXPHOS Antibody Cocktail (ab110413), STAR (ab96637), ATP assay and L-lactate assay were purchased from Abcam (Waltham, MA). PKAα/β/γ subunits antibodies were obtained from Santa-Cruz Biotechnology Inc. (Dallas, Texas) (sc-365615). siCPT1A (ON-TARGETplus Custom siRNA; cat no. CTM-3909; SMICK-000005; Dharmacon (Lafayette, CO, USA). Direct Cyclic AMP Elisa Kit was purchased from Arbor Assays (Ann Arbor, MI). HRP linked anti-mouse (#115035205) or anti-rabbit (#111035144) antibodies were obtained from Jackson ImmunoResearch Laboratories Inc. (West Grove, PA).

### Isolation and culture of bovine luteal cells

Isolation and culture of bovine luteal cells was performed as previously described (46). Bovine ovaries of early pregnant cows (fetal crown rump length < 12 cm) were collected in a local abattoir (XL Four Star Beef, Omaha, NE) and transported to the laboratory on ice in a cold M199 supplemented with antibiotics (100 U/ml penicillin-G-sodium, 100 μg/ml streptomycin sulfate, and 10 μg/ml gentamicin sulfate). The luteal tissue was dissociated with collagenase using elutriation method. The cell viability was determined by the trypan blue exclusion test, and luteal cell with viability more than 90% were used for further experiments. Enriched bovine steroidogenic small and large luteal cells (SLC and LLC, respectively) were cultured overnight on plates in basal medium (M199 containing 0.1% BSA and antibiotics) with 5% FBS at 37°C in a humidified atmosphere of 5% CO_2_. On the day of experiment, medium was removed, cells were washed with warm PBS and then fresh FBS-free medium was added, while cells were equilibrated for 2 h before applying various treatments accordingly to information included into the figure legends.

### Metabolic analysis

Sample preparation, metabolite extraction and metabolite detection were performed by the Metabolon company (Metabolon Inc., Durham, NC) based on a global unbiased platform. Briefly, the samples were prepared by using Hamilton MicroLab STAR system and the resulting extract was divided and prepared by using TurboVap (Zymark, Palo Alto, CA) and vacuum drying machine for the following instrument: i) Ultra-Performance Liquid Chromatography-Tandem Mass Spectrometry (UPLC-MS/MS) with positive ion mode electrospray ionization (Waters Acquity UPLC and Thermo-Finnigan LTQ mass spectrometer; scan range, 80–1000 m/z); ii) UPLC-MS/MS with negative ion mode electrospray ionization; iii) Liquid Chromatography (LC) polar platform; iv) Gas Chromatography-Mass Spectroscopy (GC-MS, Thermo-Finnigan Trace DSQ fast-scanning single-quadrupole mass spectrometer; scan range, 50–750 m/z). Data were extracted using Metabolon’s hardware and software (74) constructed via Microsoft’s NET technologies and compounds were identified based on Metabolon library recording *m/z*, retention time/index, and chromatographic data of more than 3,300 molecules.

### NMR sample preparation and data collection for untargeted metabolomics

The dried metabolite extracts were reconstituted in 600 µL of 50 mM phosphate buffer in 99.8% D_2_O (Isotec, St. Louis, MO) at pH 7.2 (uncorrected). 50 µM of TMSP(3-(tetramethysilane) propionic acid-2,2,3,3-d4) were added for spectral referencing. The solution was mixed by gentle vortexed and centrifuged at 13000 rpm for 5 min, and then the supernatant was transferred to 5 mm NMR tube. The NMR data was collected using Bruker AVANCE III 700 MHz spectrometer equipped with 5 mm triple-resonance cryogenic probe (^1^H, ^13^C, and ^15^N) with a Z-axis gradient were used to acquire the NMR data. The sample collection was automated using a SampleJet sample changer, ATM (automatic tuning and matching), and Bruker IconNMR™ software. The NMR experiment was conducted at 300 K with 32 K data points, 256 scans, 16 dummy scans, and a spectral width of 11160 Hz using an excitation sculpting pulse sequence.

### NMR sample preparation and data collection for targeted metabolomics using stable isotope tracers

The dried metabolite extracts were reconstituted in 450 µL of 50 mM phosphate buffer in 99.8% D_2_O (Isotec, St. Louis, MO) at pH 7.2 (uncorrected). 500 µM of TMSP(3-(tetramethysilane) propionic acid-2,2,3,3-d4) were added for spectral referencing. The solution was mixed by gentle vortexed and centrifuged at 13000 rpm for 5 min, and then the supernatant was transferred to 5 mm NMR tube. The NMR data was collected using Bruker AVANCE III 700 MHz spectrometer equipped with 5 mm cryoprobe. The sample collection was automated using SampleJet and Bruker IconNMR™ software. A 2D ^1^H-^13^C Heteronuclear Single Quantum Coherence (HSQC) spectra were collected at 300 K with 2 K data points in the direct dimension, and 256 data points in the indirect dimension for 24 scans, 16 dummy scans.

### NMR data analysis for untargeted and targeted metabolomics

MVAPACK software (http://bionmr.unl.edu/mvapack.php) were used to analyze the untargeted metabolomics NMR data (75) Principal Component Analysis (PCA) model was generated using raw NMR data that was Fourier transformed, automatically phased, and binned using adaptive intelligent binning, normalized using standard normal variate normalization, and noise was removed as stated by (76–79). Orthogonal Projections to Latent Structures Discriminant Analysis (OPLS-DA) scores and back-scaled loadings were generated using full intact spectrum. Spectra were normalized using standard normal variate normalization, noise was removed as stated earlier and scaled using Pareto scaling. The Human Metabolomics Database and Chenomx NMR Suite 7.6 (http://www.chenomx.com/) were used for metabolite identification(78, 80)(78, 80)(78, 80)(77, 79)(76, 78)(76, 78)(76, 78)(77, 79)(78, 80)(78, 80)(78, 80)(78, 80)(78, 80)(78, 80)(78, 80)(79, 81)(79, 81)(79, 81)(78, 80)(79, 81)(78, 80)(77, 79). 2D ^1^H-^15^N HSQC spectra were processed using NMRPipe, and the time series was analysed using NMRviewJ to obtain peak intensities over the course of the experiment, as previously explained.

### Seahorse analysis

*Mito Stress Assay:* Mitochondrial oxygen consumption rate (OCR) was measured using a Seahorse XF^e^96 analyzer (Seahorse Bioscience, North Billerica, MA, USA). In brief, primary luteal cells were plated on XF^e^96 cell culture microplate (40×10^4^ cells per well) and incubated overnight. Next day, cells were washed with warm PBS and medium was replaced with fresh one. Cells were pre-treated for 60 min with inhibitors, either Etomoxir (30 µM) or UK5099 (1 μM). Then, medium was changed with assay medium (base medium with L-glutamine, glucose, and pyruvate) and cells were incubated with LH (10 ng/ml) for 60 min in non-CO_2_ incubator. The assay was done following injections: oligomycin (2 µM), FCCP (2-(2-(4-(trifluoromethoxy) phenyl) hydrazinylidene) - propanedinitrile) (0.5 µM), and rotenone (0.5 µM) combined with antimycin (AA; 0.5 µM). Before and after each injection were performed three measurements of OCR and extracellular acidification rate (ECAR) for each 3 minutes. First, the basal oxygen consumption rate (basal respiration) was measured. Afterwards, oligomycin, which inhibits ATP synthase activity, was injected to reveal ATP production coupled with mitochondria respiration. Next, FCCP that increases proton pump was injected and allowed maximal mitochondrial respiration. Finally, rotenone combined with antimycin A was injected to inhibit the flux of electrons through complexes I and III; so, the remaining oxygen consumption rate was primarily due to non-mitochondrial respiration. The spare respiratory capacity was calculated by subtracting the basal respiration from the maximal respiration. At least four technical replicates were used for each treatment, and the experiments were repeated three times to confirm the results.

#### Glycolysis Stress Assay

The Glycolysis Stress Kit was applied to determine glycolysis in luteal cells. Similarly, primary luteal cells were plated on XF^e^96 cell culture microplate (40×10^4^ cells per well) and incubated overnight. Next day, cells were washed with warm PBS and medium was replaced with fresh one. Before assay, cells were pretreated with either Shikonin (5 µM) or UK5099 (1 μM) in culture medium. Afterwards, cultured medium was replaced with assay medium (base medium supplemented with 2 mM L-glutamine) and treated with LH (10 ng/ml) in non-CO_2_ incubator. The following injections were performed during the assay: glucose (10 mM), oligomycin (2 μM) and 2-DG (5 mM). Before and after each injection, OCR and ECAR were measured three times each three minutes. Four technical and biological replicates were used for each treatment group.

### Lactate concertation assay

To confirm glycolysis, media samples were collected after Seahorse analysis to determine lactate concentration using the L-lactate Assay according to manufacturer’s protocol (Abcam). Absorbance values at 450 nm were obtained using a plate reader. Readings were corrected for background absorbance using the absorbance value from the assay medium only, and the corrected values were applied to a standard curve to calculate extracellular lactate levels.

### Glucose uptake assay

Luteal cells (10,000 per well) were plated on 96-well plates and cultured overnight in M199 supplemented with antibiotics and 5% FBS. Next day, media was removed, cells were washed with PBS and fresh media without FBS was added. After two hours of equilibration, cells were treated with LH (10 ng/ml) for 10 and 60 min. Afterward, media was removed, cells were washed with PBS and 1 mM 2-deoxy-D-glucose was added. After 10 min of incubation at room temperature, the stop buffer followed by the neutralization buffer were added and the cells were briefly shaken. Then, 2-deoxy-D-glucsoe-6-phopshate detection reagent was added and cells were incubated for 45 min at room temperature. Luminescence was recorded using luminometer FluroStar Optima (BMG Labtech).

### ATP Assay

Luteal cells were plated on 96-well plates with clear bottom and cultured overnight in M199 supplemented with antibiotics and 5% FBS. Next day, media was removed, cells were washed with PBS and fresh media without FBS was added. After two hours of equilibration, cells were pretreated with inhibitors BMS303141 (10-50 µm) or Teglicar (10-50 µM) for 60 min and then treated with LH (10 ng/ml) for 240 min. Afterward, ATP assay was performed according to manufacturer’s protocol. Cells were lysed for 5 min by using detergent solution supplied in the assay, and then substrate solution was added, and cells were incubated for 5 min, adapted in dark for 10 min and luminescence was measured by using luminometer FluroStar Optima (BMG Labtech, Cary, NC).

### cAMP assay

cAMP concentration was measured in media samples collected after incubation of luteal cells with inhibitor and/or LH. Standards and media samples were pipetted onto 96-well coated with an antibody to capture sheep IgG following primer plate. Next, a cAMP-peroxidase conjugate was added to both standards and samples. After 2h incubation, plate was washed, and substrate was added. After short incubation, the reaction was stopped, and the intensity of the generated color was detected in a microtiter plate reader at 450nm. Data were analyzed by using online tool provided by manufacturer at MyAssays website (www.myassays.com/arbor-assays-cyclic-amp-direct-eia-kit-non-acetyl.assay).

### Western Blot Analysis

After *in vitro* experiments cells were collected and lysed for Western blot analysis as previously described (46). Lysates were subjected to separation on 10% SDS-PAGE and transferred onto nitrocellulose membranes, which were then blocked with 5% BSA in Tris-buffered saline with 0.1% Tween-20 (TBST) at room temperature for 1 h. Then, membranes were incubated with primary antibodies at 4°C overnight. Next day, membranes were washed three times with TBST and blocked in secondary anti-mouse or anti-rabbit HRP-conjugated antibodies (1: 10,000) in 5% non-fat milk in TBST for 1 h, followed by a second series of three washes in TBST. Blots were imaged and quantified using iBright CL1500 Imaging System (Thermo Fisher Scientific Inc., Waltham, MA).

### siRNA transfection

*CPT1A* was knocked down using specific CPT1A silencing RNA (siRNA) to determine the role of CPT1A on luteal progesterone production, cAMP production, PKA signaling and steroidogenic machinery or mitochondrial proteins. Small luteal cell populations were transfected with Lipofectamine RNAimax and control siRNA (siCTL) or si*CPT1A* (ON-TARGETplus Custom siRNA (CTM-385778; HOUSF-000005) in opti-MEM culture medium. After 6 hours, 5% FBS was added to culture media and cells were maintained for 48 hours. Knockdown of *CPT1A* was confirmed by western blotting. Following transfection, luteal cells were treated with LH (10 ng/mL) for 4 hours. Media were collected for progesterone concentration analysis and cell lysates were immediately collected for further western blotting analysis.

### Progesterone ELISA

Luteal cells were cultured in 48-well plates. Media samples were collected to determine progesterone concentration using ELISA method (Diagnostic Systems Laboratories, Inc., Webster, TX.) according to the manufacturer’s instructions.

### Statistical analysis

Statistical analyses were conducted using GraphPad Prism v. 8.0 software (GraphPad Software, Inc., San Diego, CA, USA). All experiments were performed at least three times using luteal cells obtained from different animals. The data are presented as the means ± SEM of the averages from multiple experiments. Data were analyzed by one- or two-way ANOVA, followed by Bonferroni multiple comparison tests. Statistical differences were considered as significant at P < 0.05.

---

**Supplementary Figure 1.**
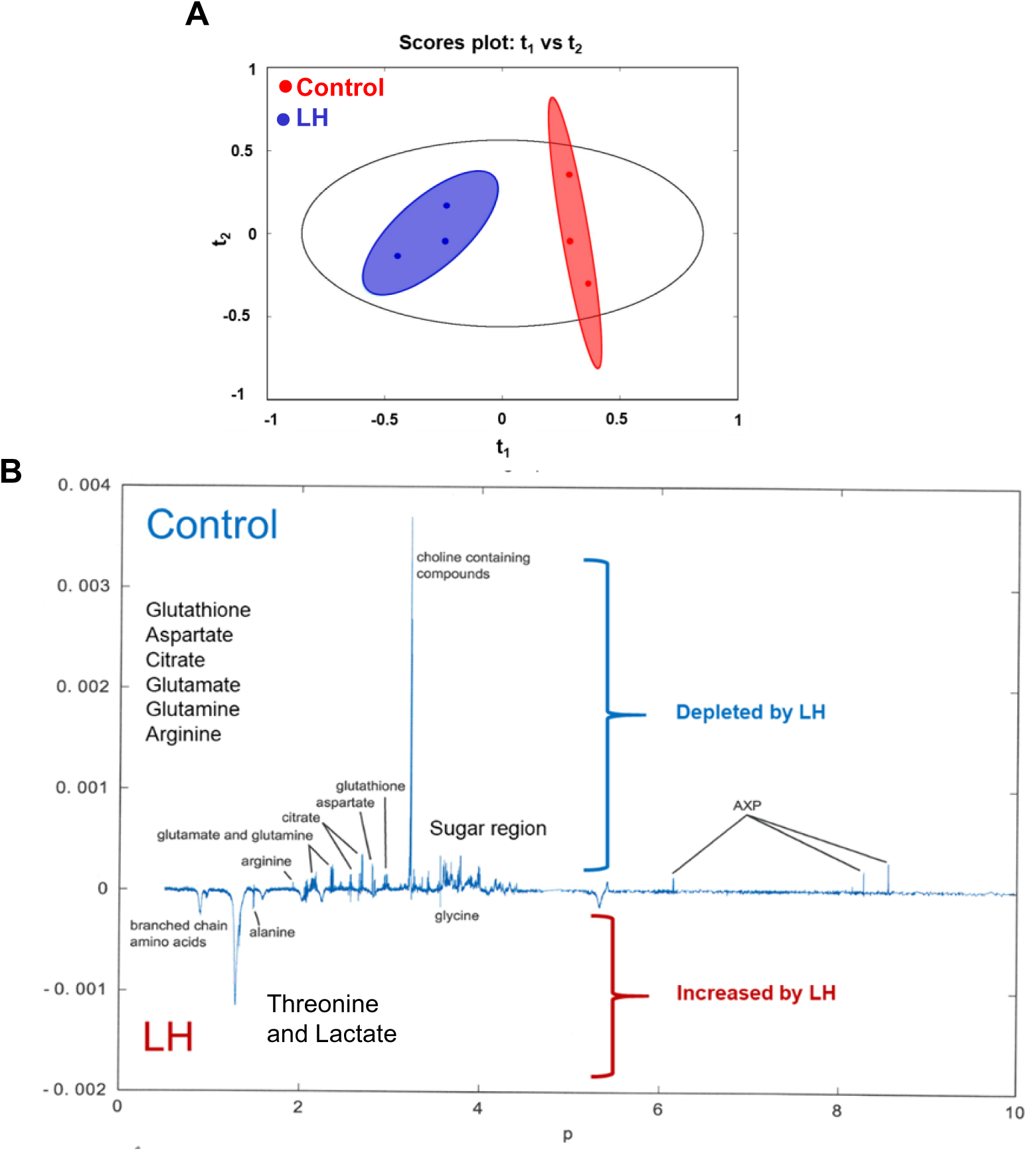
Enriched preparation of small luteal cells treated with LH (10 ng/ml) for 240 min were analyzed by Nuclear Magnetic Resonance (NMR) spectroscopy. (**A**) Principal Component Analysis (PCA) plot showing grouping of sample preparation (n=3 per group) from untreated (Control; red circle) and LH-treated cells (blue circle). (**B**) NMR spectrum showing the most significantly (depleted or increased) changed metabolites post-LH treatment.

**Supplementary Figure 2.**
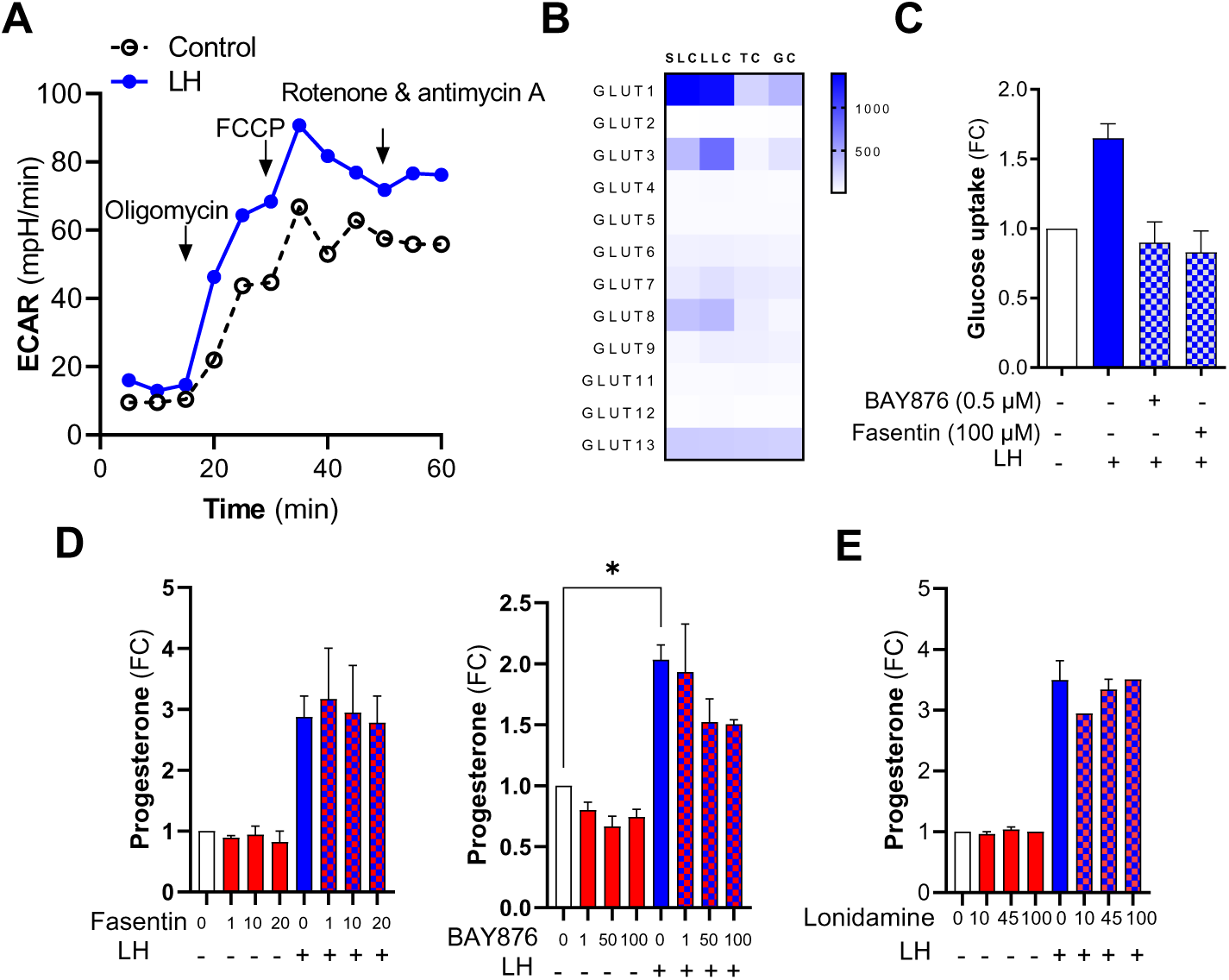
(**A**) Representative graph showing changes in the extracellular acidification rate (ECAR) in untreated (Control) and LH-treated cells (60 min) based on Seahorse Cell Mito Stress Test. (**B**) Heatmap representing expression of glucose transporters (*GLUT 1-9*; *GLUT 11-13*) in the small and large luteal cells (SLC and LLC) as well as theca and granulosa cells (TC and GC). Data were obtained from previously performed microarray analysis (GSE83524). (**C**) Glucose uptake measured by using luminescence method in the small luteal cells pretreated with inhibitors of GLUT1 (BAY876; 1 µM) or GLUT1/4 (Fasentin; 100 µM) and then treated with LH (10 ng/ml) for 60 min. Data are presented as a fold change (FC) and mean±SEM (n=2). (**D-E**) Progesterone production by the small luteal cells pretreated with inhibitor of Glut1 (BAY876; 1-100 μM) or Glut1/4 (Fasentin; 1-20 μM) or hexokinase (Lonidamine; 10-100 μM) and then treated with LH (10 ng/ml) for 240 min. Data are presented as a fold change (FC) and mean±SEM (n=2-3).

**Supplementary Figure 3.**
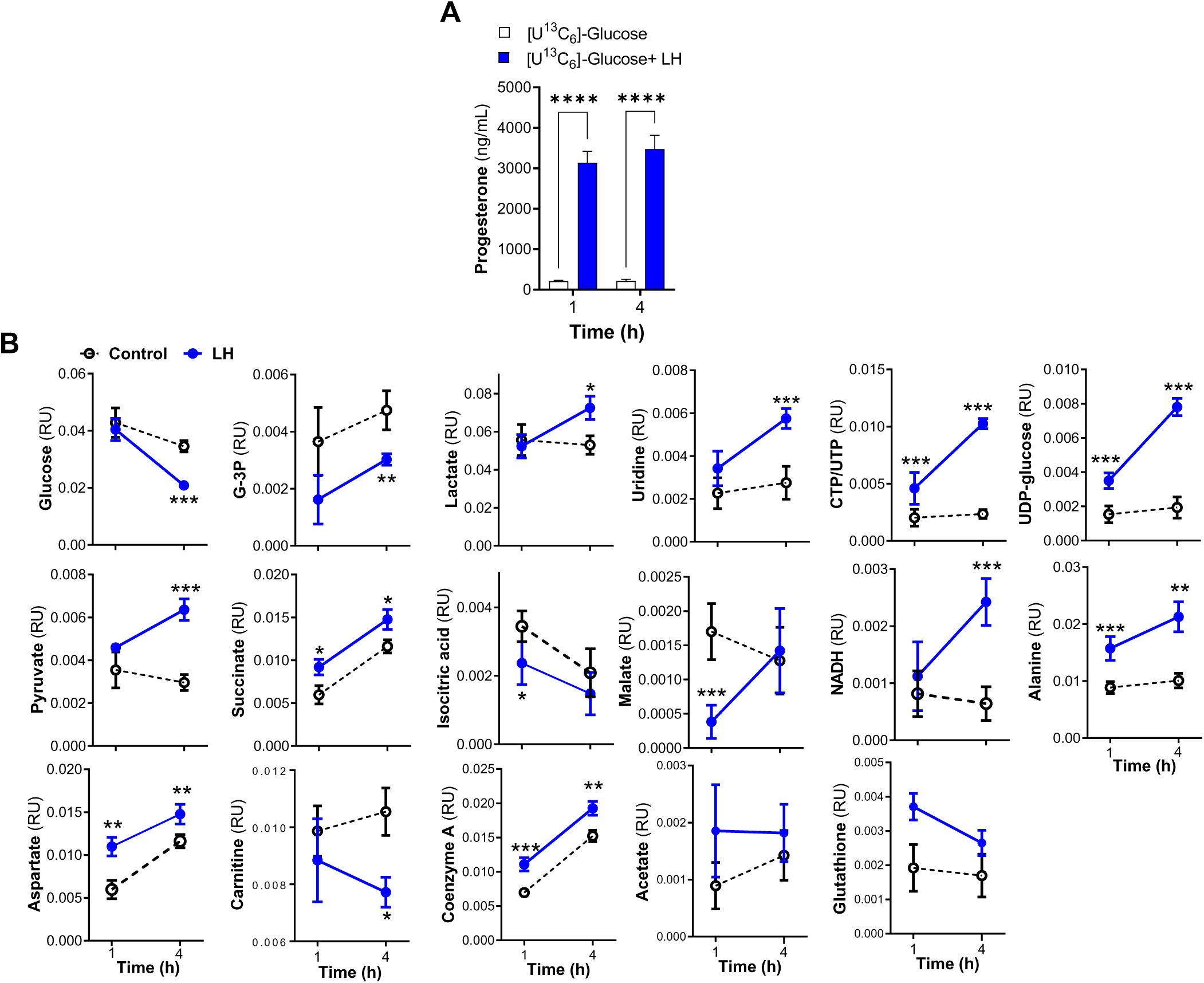
(**A**) Progesterone production by the small luteal cells treated with [U^13^C_6_]-labeled glucose (5 mM) alone or in the presence of LH (10 ng/ml) for 60- and 240-min. Data are represented as mean±SEM (n=6) and were analyzed using one-way ANOVA test. Asterisks **** mean significant change with P < 0.0001. (**B**) Time-dependent changes in the concentration of selected metabolites in small luteal cells cultured in medium with [U-^13^C_6_]-labeled glucose in the presence or absence (Control; black dotted line) of LH (blue solid line). Data are presented as relative units (RU) and mean±SEM (n=6). Asterisks *, **, *** mean significant change with P < 0.05, P < 0.01, and P < 0.001, respectively.

**Supplementary Figure 4.**
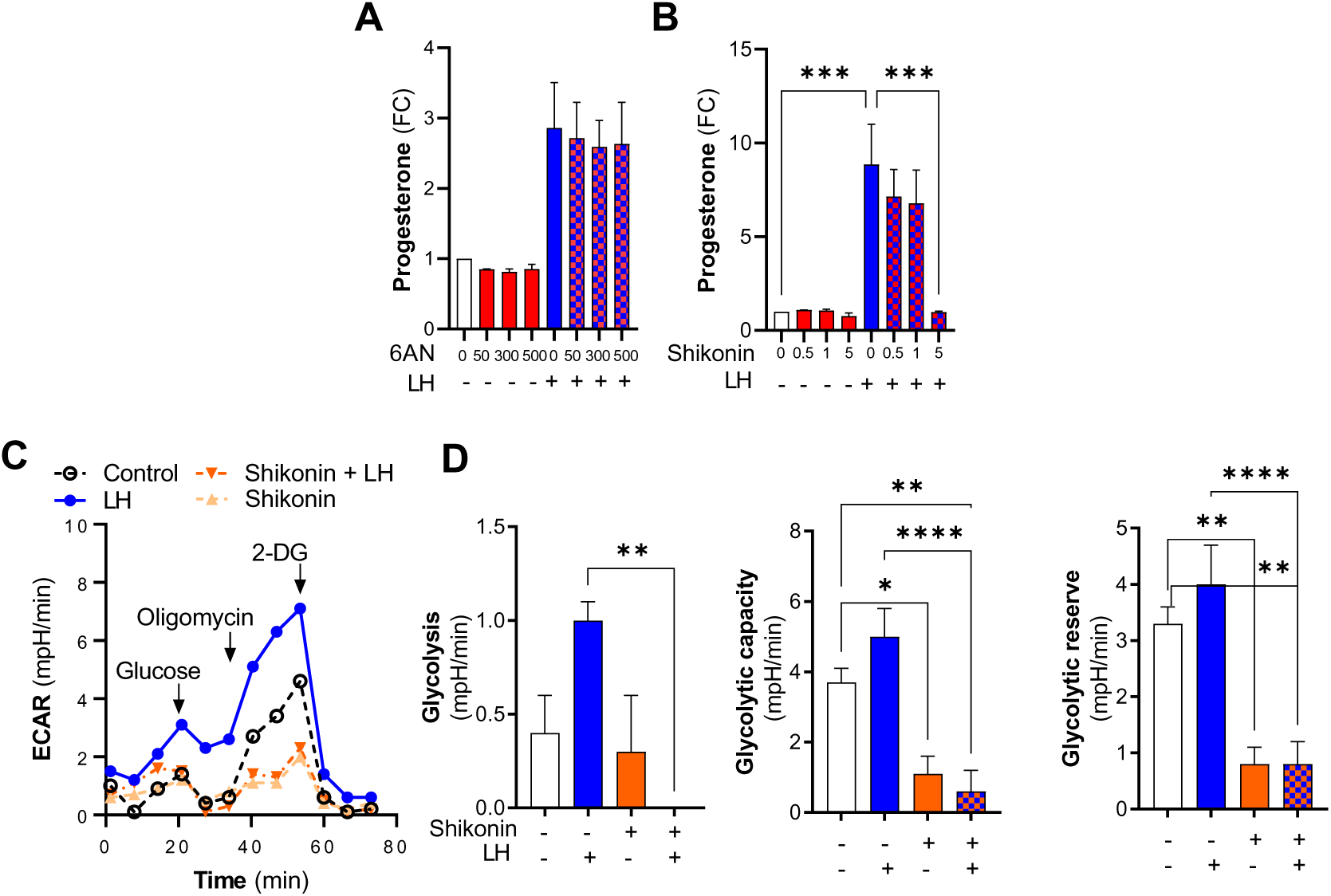
(**A-B**) Progesterone production by cells pretreated with inhibitor of glucose-6-phosphate dehydrogenase (G6PD; 6AN; 50-500 μM) or pyruvate kinase (Shikonin; 0.5-5 μM) and then treated with LH (10 ng/ml) for 240-min. Data represented as fold change (FC) and mean ± SEM (n=3-4) were analyzed using one-way ANOVA test. Asterisks *** mean significant change with P < 0.001. (**C**) Glycolytic rate was analyzed using Seahorse Glycolytic Rate Assay. Representative graph showing extracellular acidification rate (ECAR) in the small luteal cells pretreated with pyruvate kinase inhibitor (Shikonin) and then treated with LH (10 ng/ml) for 60 min. (**D**) Glycolysis, glycolytic capacity, and glycolytic reserve in the small luteal cells pretreated with pyruvate kinase inhibitor (Shikonin) and then treated with LH. Data are represented as mean ± SEM (n=10-12) and were analyzed using a one-way ANOVA test. Asterisks *, ** and **** indicate significant change with P < 0.05, P< 0.01 and P < 0.0001, respectively.

**Supplementary Figure 5.**
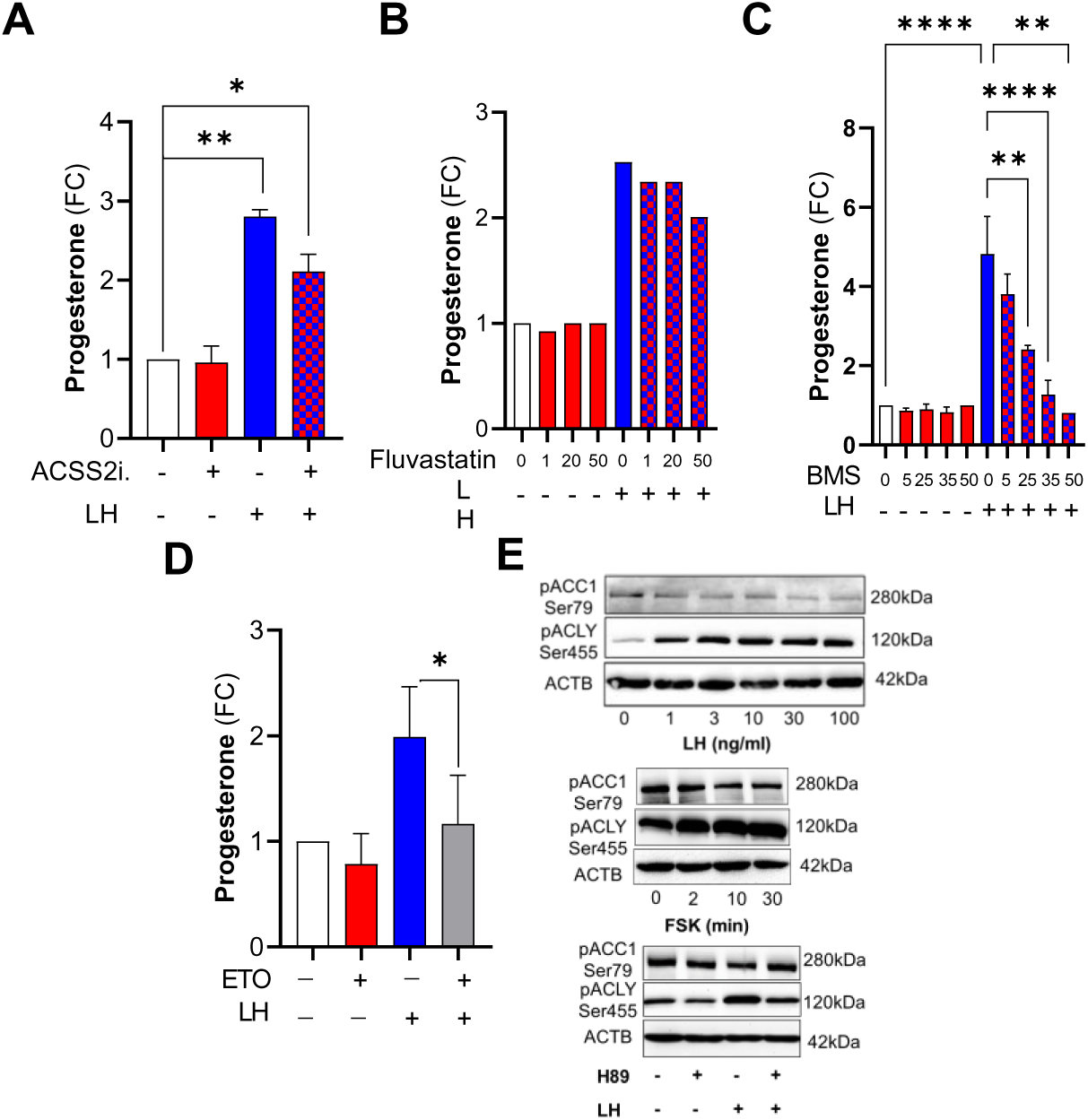
(**A-D**) Progesterone production by the small luteal cells pretreated with inhibitor of ACSS2 (ACSS2 i.; 10 µM), HMGCR (Fluvastatin; 1-50 µM), ACLY (BMS303141; 5-50 µM) or Etomoxir for 60 min and then treated with LH (10 ng/ml) for 240 min. Data are presented as a fold change (FC) and mean±SEM (n=2-5). Data were analyzed using a one-way ANOVA test. Asterisks *, ** and **** mean significant change with P < 0.05, P < 0.01 and P < 0.0001, respectively. (**E**) Representative blots showing phosphorylation of ACLY Ser455 and ACACA Ser89 in the small luteal cells treated with LH (1-100 ng/ml) for 30 min or cAMP/PKA activator forskolin (FSK; 10 μM) for 2-30 min as well as cells pretreated with PKA inhibitor (H89; 20 µM) and then treated with LH (10 ng/ml) for 10 min.

**Supplementary Figure 6.**
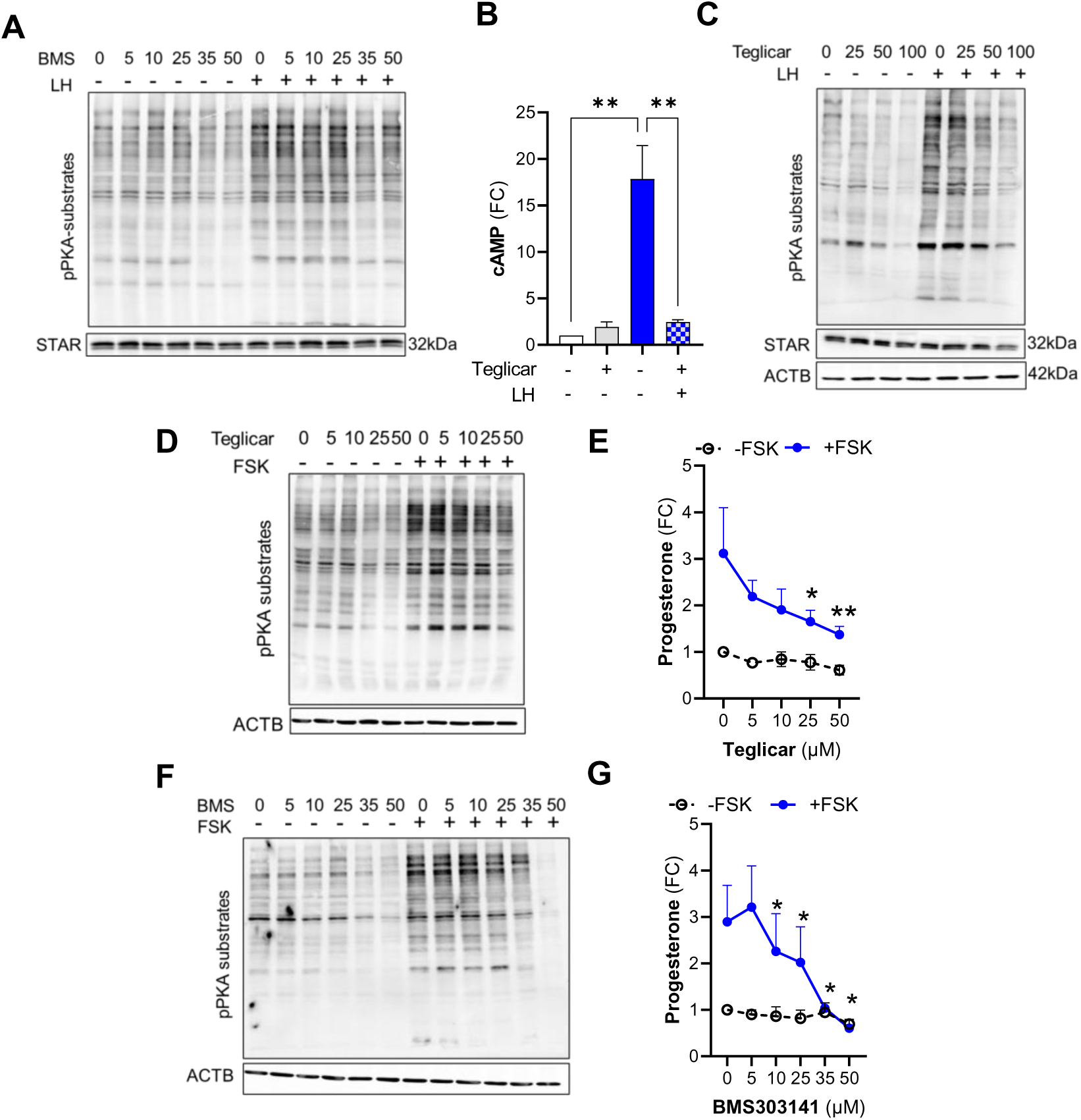
(**A**) Representative blots showing phosphorylation of PKA substrates and content of STAR in the small luteal cells pretreated with ACLY inhibitor (BMS303141; 5-50 µm) and then treated with LH (10 ng/ml) for 240 min. (**B**) cyclic AMP (cAMP) production by small luteal cells pretreated with CPT1A inhibitor (Teglicar; 25 µM) and then treated with LH (10 ng/ml) for 240 min. Data are presented as a fold change (FC) and mean±SEM (n=2-5). Data were analyzed using a one-way ANOVA test. Asterisks ** mean significant change with P < 0.01. (**C-D**) Representative blots showing phosphorylation of PKA substrates and content of STAR in the small luteal cells pretreated with CPT1A inhibitor (Teglicar; 5-50 µm) and then treated with LH (10 ng/ml) or PKA activator Forskolin (FSK; 10 µM) for 240 min; (**E**) Progesterone production by small luteal cells pretreated with Teglicar (5-50 µM) and then treated with PKA activator Forskolin (FSK; 10 µM) for 240 min. (**F**) Small luteal cells were pretreated with inhibitor of ACLY (BMS303141; 5-50 µM) and then treated with PKA activator forskolin (FSK; 10 µM) for 240 min. Representative blots of phosphorylation of PKA substrates in the small luteal cells pretreated with ACLY inhibitor (BMS303141) and then treated with forskolin (FSK). (**G**) Progesterone production by the small luteal cells pretreated with ACLY inhibitor (BMS303141) and then treated with PKA activator forskolin (FSK). Data are presented as a fold change (FC) and mean±SEM (n=2-5). Data were analyzed using a two-way ANOVA test. Asterisks *, P < 0.05.

